# Antiviral effects of miRNAs in extracellular vesicles against severe acute respiratory syndrome coronavirus 2 (SARS-CoV-2) and mutations in SARS-CoV-2 RNA virus

**DOI:** 10.1101/2020.07.27.190561

**Authors:** Jae Hyun Park, Yuri Choi, Chul-Woo Lim, Ji-Min Park, Shin-Hye Yu, Yujin Kim, Hae Jung Han, Chun-Hyung Kim, Young-Sook Song, Chul Kim, Jisook Moon

## Abstract

Severe acute respiratory syndrome coronavirus 2 (SARS-CoV-2) causes coronavirus 2019 (COVID-19). No treatment is available. Micro-RNAs (miRNAs) in mesenchymal stem cell-derived extracellular vesicles (MSC-EVs) are potential novel therapeutic agents because of their ability to regulate gene expression by inhibiting mRNA. Thus, they may degrade the RNA genome of SARS-CoV-2. EVs can transfer miRNAs to recipient cells and regulate conditions within them. MSC-EVs harbor major therapeutic miRNAs that play important roles in the biological functions of virus-infected host cells. Here, we examined their potential impact on viral and immune responses. MSC-EVs contained 18 miRNAs predicted to interact directly with the 3’ UTR of SARS-CoV-2. These EVs suppressed SARS-CoV-2 replication in Vero E6 cells. In addition, five major miRNAs suppressed virus activity in a luciferase reporter assay by binding the 3’ UTR. MSC-EVs showed strong regenerative effects and potent anti-inflammatory activity which may prevent lethal cytokine storms. We confirmed that EVs regulated inflammatory responses by several cell types, including human brain cells that express the viral receptor ACE2, suggesting that the brain may be targeted by SARS-CoV-2. miRNAs in MSC-EVs have several advantages as therapeutic agents against SARS-CoV-2: 1) they bind specifically to the viral 3’ UTR, and are thus unlikely to have side effects; 2) because the 3’ UTR is highly conserved and rarely mutates, MSC-EV miRNAs could be used against novel variants arising during viral replication; and 3) unique cargoes carried by MSC-EVs can have diverse effects, such as regenerating damaged tissue and regulating immunity.

## Introduction

Severe acute respiratory syndrome coronavirus 2 (SARS-CoV-2) is the positive sense single-stranded RNA virus that causes coronavirus 2019 (COVID-19). SARS-CoV-2 is a variant of the coronavirus SARS-CoV, which is associated with severe acute respiratory syndrome. SARS-CoV-2 is the seventh coronavirus known to infect humans; the others are 229E, NL63, OC43, HKU1, MERS-CoV, and the original SARS-CoV. SARS-CoV-2 is a Sarbeco virus (β-CoV strain B) with an RNA genome of about 30,000 bases. The virus attaches to the angiotensin converting enzyme 2β (ACE2) receptor on human cells. After binding to a host cell, the cellular protease TMPRSS2 cleaves the viral spike protein to release the fusion peptide. Then the virion releases RNA into the cell, leading to virus replication and infection of other cells. Since SARS-CoV-2 was identified as the causative agent of COVID-19, scientists have sought to understand the genetic composition of the virus and to find ways to effectively treat the infection. No specific treatments for COVID-19 have been developed. Various treatments have been proposed, and some approved drugs appear to be associated with positive results; however, more work is required before a viable treatment can be deployed in the clinic. Developing new vaccines takes time, and clinical trials require rigorous testing and safety tests. The RNA genome has a high rate of mutation, making RNA viruses very heterogeneous [1]. The mutation rate of viral RNA is about 1 million times greater than that of human DNA [2]. Consequently, RNA viruses show large genetic diversity, a property important for their survival [3, 4]. The high frequency of viral gene mutations makes it difficult to develop therapeutic agents. For example, antibody-based therapeutics developed against one variant must be redesigned after the virus mutates. Consequently, influenza vaccines need to be updated annually. Also, viral variants develop resistance to antiviral drugs, and mild virus variants become virulent spontaneously. Consequently, therapeutic and preventive agents that can cope with viral mutations are needed urgently.

MicroRNAs (miRNAs) are small non-coding RNAs of 18–25 nucleotides (nts); these small molecules are important regulators of gene expression because they suppress messenger (m)RNAs. miRNAs regulate approximately 30–70% of human gene expression by matching their “seed” sequence with complementary mRNA [5]. Alongside the regulation of human gene expression, miRNAs bind directly to viral RNA genomes or affect viral replication by mediating changes in the host genome. Several miRNAs bind directly to a wide range of RNA viruses and modulate pathogenesis [6]. In addition, the seed sequence, which is located at the 5’ end of the miRNA, binds to a complementary sequence within the 3’ UTR of the invading viral RNA [7]. A perfect match between miRNA and the whole mRNA target sequence leads to RNA cleavage, although this is rare in mammals [7]. More often, exact seed sequence matching results in translational inhibition, followed by RNA degradation [8, 9]. Only seven nucleotides within a miRNA sequence are required to bind target mRNAs because either partial or perfect binding between miRNA and mRNA results in the degradation of the target mRNA strand [10, 11]. EVs are small membrane-bound bodies that contain cargoes such as nucleic acids, proteins, lipids, and metabolites [12]. Release of EVs is an important form of intercellular communication that plays roles in the physiological and pathological processes underlying multiple diseases. One of the most fascinating features of EVs is their ability to transfer miRNAs to recipient cells and regulate cell conditions [13-15]. For example, miRNAs delivered by EVs to both immune cells and other cell types repress or degrade target mRNAs in recipient cells [16-18].

The regulatory or regenerative functions of miRNAs contained in stem cell-derived EVs have attracted attention as a novel therapeutic approach [19]. EVs secreted from mesenchymal stem cells (MSCs) mediate tissue regeneration and help to repair damage in a variety of diseases, including cardiac ischemia, liver fibrosis, and cerebrovascular disease. Multiple studies report the therapeutic effects of miRNAs delivered to host cells by MSC-EVs. MSC-EVs transport proteins, ncRNA, miRNAs, and lipids, which mediate cardiac tissue repair and regulate the environment around damaged tissue, induce angiogenesis, promote proliferation, and prevent apoptosis [20, 21]. When SARS-CoV-2 attaches to the ACE2 receptor to invade a host cell, immune cells such as T lymphocytes and dendritic cells (DCs) secrete and absorb EVs containing miRNAs that attack the infected viral RNA. Thus, delivery of miRNA by MSC-EVs represents a new therapeutic mechanism for combating SARS-CoV-2 and regulating the pro-inflammatory environment.

Here, we identified candidate therapeutic miRNAs with important roles in the biological functions of virus-infected host cells, and characterized the antiviral effects of miRNAs derived from placenta-derived MSC-EVs (pMSC-EVs) or placenta EVs, which exhibit potent regenerative and anti-inflammatory effects. Five selected miRNAs (miR-92a-3p, miR-103a-3p, miR-181a-5p, miR-26a-5p, and miR-23a-3p) blocked SARS-CoV-2 RNA replication and suppressed virus-mediated pro-inflammatory responses by human bronchial epithelial cells and lung fibroblasts, all of which express ACE2 receptors. These effects were also observed in human brain cells and brain immune cells. Interestingly, we found that ACE2 is expressed in human fetal brain stem cells, suggesting that the brain could be target of SARS-CoV-2. Most importantly, we found that the five miRNAs bound to the 3’ UTR of SARS-CoV-2, the sequence of which is conserved among coronaviruses.

## Results

Antiviral effect of EVs against SARS-CoV-2Viral infections cause a variety of cellular reactions, and viruses themselves are known to create an environment that allows for replication and spread. In addition, the virus causes apoptosis, and the completed virus is easily transmitted. In addition, host cells are known to secrete various cytokines and stimulate immune cells by their protective action. Consequently, it is very important to break down viral replication before stimulation of inflammatory cytokines. Figure 1 shows the hypothesis, which asserts the antiviral effect of miRNA in EVs and EVs against SARS-CoV-2, and explains how miRNA directly degrades viruses and indirectly suppresses excessive immune responses. SARS-CoV-2 viruses invade cells through ACE2 receptors. They penetrate cells by binding to these receptors and transporting viral RNAs into the cytoplasm where they replicate. When the cell recognizes viral RNAs in the cytoplasm, it secretes cytokines, induces an immune fraction, and stimulates immune cells to fight the viral infection. However, if too much cytokine is secreted due to excessive stimulation, the immune cells are over-activated and can, damage normal cells and tissues. Resulting in a phenomenon is referred to as a cytokine storm. EVs enter cells through various pathways, including membrane fusion, receptor-mediated uptake, and active endocytosis. We demonstrated that key miRNAs expressed in MSC-EVs degrade SARS-CoV-2 RNAs by interacting directly with the 3’ UTR. In addition, the miRNAs in EVs exerted an anti-inflammatory effect, which prevented the cytokine storms by dampening the excessive immune response caused by the virus (**Fig. 1, left panel**). Specifically, the direct effects of EV miRNAs against SARS-CoV-2 virus regulation are mediated by targeting regions within the SARS-CoV-2 genome, including the 3’ UTR, the 5’ UTR, and coding sequences. Particularly, direct binding to the 3’ UTR is predicted to down-regulate SARS-CoV-2 RNA. In addition, EVs regenerate damaged tissue and regulate the pro-inflammatory environment via their miRNAs and protein cargoes, indicating their potential to suppress cytokine storms caused by viral infection (**Fig. 1, right panel**). Cargoes including miRNAs in MSC-EVs attenuate induced inflammation and apoptosis caused by SARS-CoV-2, and suppress the expression of transcription/translation machinery involved in virus replication and translation, thereby indirectly suppressing the action of virus.

**Fig. 1.**
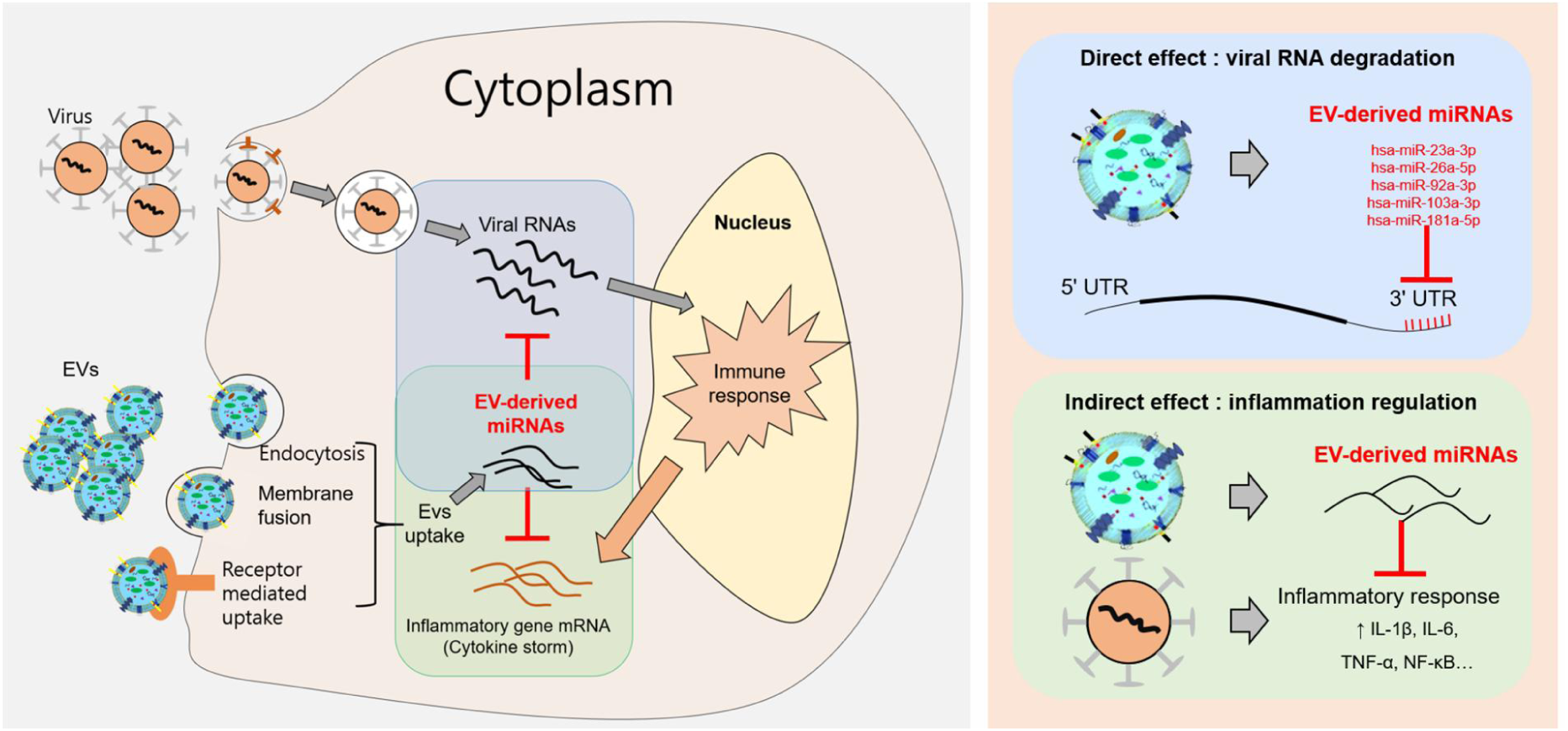
Mechanism underlying the antiviral effects of MSC-EV-derived miRNAs against SARS-CoV-2. Our hypothesis of antiviral effects of miRNAs in EVs and EVs against SARS-CoV-2. SARS-CoV-2 viruses invade cells through ACE2 receptors and EVs enter cells through various pathways, including membrane fusion, receptor-mediated uptake, and active endocytosis. The cell secretes cytokines, induces an immune fraction, and stimulates immune cells to fight the viral infection. Too much cytokine leads to a cytokine storm (left panel). The direct effects of EV miRNAs on SARS-CoV-2 virus regulation are mediated by targeting regions within the SARS-CoV-2 genome, including the 3’ UTR, the 5’ UTR, and coding sequences. In particular, direct binding to the 3’ UTR is predicted to down-regulate SARS-CoV-2 RNA. In addition, EVs regenerate damaged tissue and regulate the proinflammatory environment via their miRNAs and protein cargoes, indicating their potential to suppress cytokine storms caused by viral infection (right panel).

EVs isolated from placental stem cells (pMSCs) were positive for the common EV marker CD63 (**Suppl. Fig. 1A**). The hydrodynamic diameter of EVs measured by nanoparticle tracking analysis (NTA) was 121.8 nm (**Suppl. Fig. 1B**) and a representative transmission electron microscopy (TEM) image exhibited the typical EV morphology (**Suppl. Fig. 1C**). In addition, Western blotting detected typical EV markers, including CD81, CD63, TSG101, and CD9 (**Suppl. Fig. 1D**). To verify the effect of EVs against SARS-CoV-2, we first examined the cytotoxicity of EVs in an MTT assay using Vero cells. Vero cells were treated for 24 h with two-fold serial dilutions of EVs (0.003-4.7 μg). No cytotoxicity was observed at any concentration tested and rather cells proliferated after treatment with EVs at all doses (**Fig. 2A**, *Ps* < 0.05). Subsequently, we developed a 96-well plate assay in which live cells were stained with crystal violet, whereas cells showing virus-mediated Cytopathic Effect (CPE) were not. To confirm the cytotoxic concentration of SARS-CoV-2, Vero cells were infected with the virus (10-fold serial dilutions: 10^1^ TCID_50_/well, 10^2^ TCID_50_/well, and 10^3^ TCID_50_/well, n=3 replicates) and incubated for 3 days. The amount of cell death after 3 days was confirmed by examining CPE. Cell detachment was observed at doses of 10^1^ TCID_50_/well to 10^3^ TCID_50_/well (**Fig. 2B, left panel**). Next, we examined the antiviral effects of EVs against SARS-CoV-2 at three different TCID_50_ concentrations. EVs at 5 μg, 2.5 μg, and 1.25 μg showed 100%, 100%, and 66% efficacy, respectively, at the titer of 10^1^ TCID_50_. EVs at 5 μg and 2.5 μg showed 66% and 33% efficacy, respectively, at a titer of 10^2^ TCID_50_. However, EVs did not show antiviral activity at a virus titer of 10^3^ TCID_50_ (**Fig. 2B, right panel**).

**Fig. 2.**
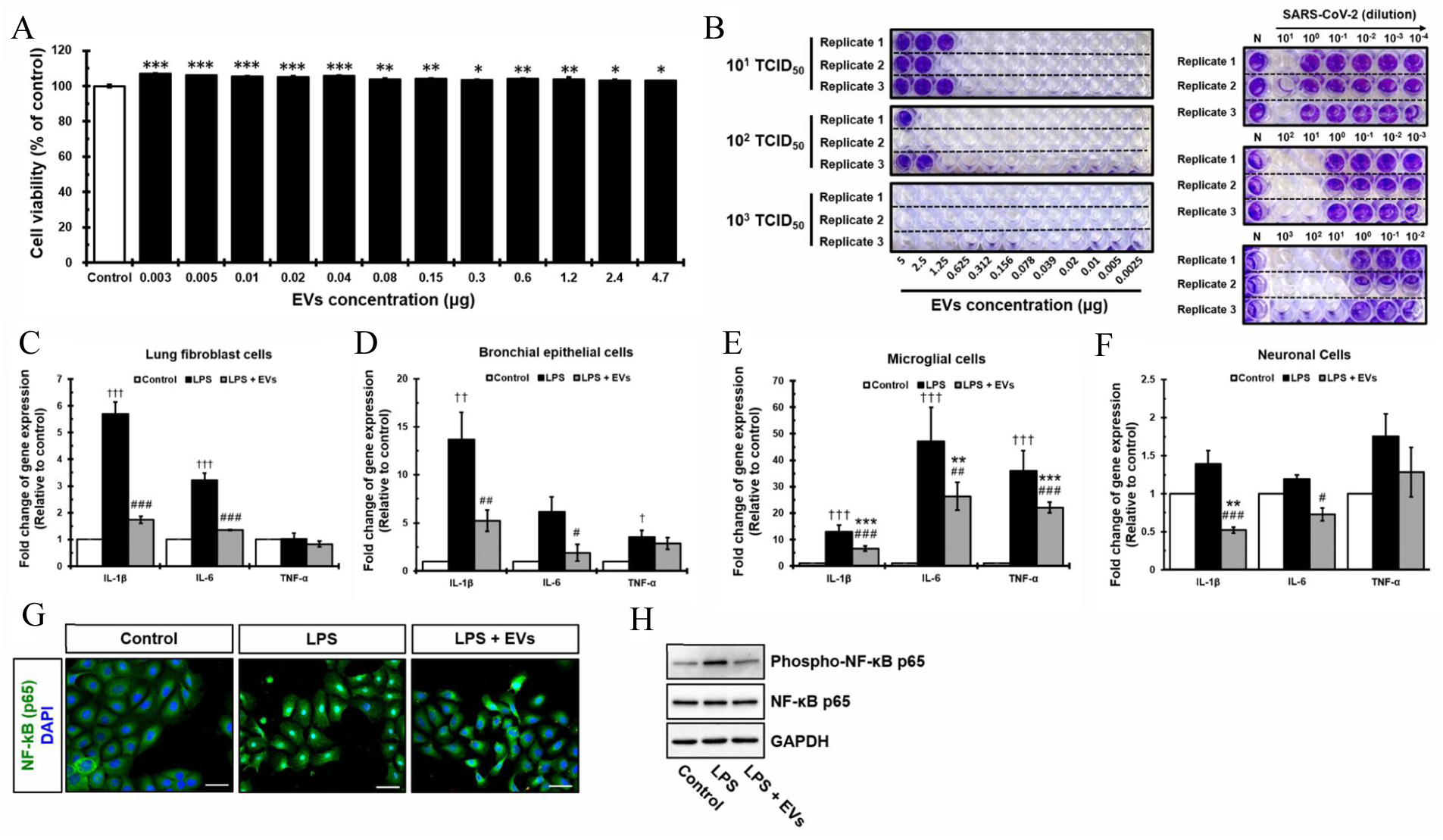
Antiviral effect of EVs against SARS-CoV-2. **(A)** The cytotoxicity of EVs evaluated by using the MTT assay on Vero cells. (**B)** The anti-SARS-CoV-2 activity of EVs was tested using established cell-based screening assay. Vero cells were infected with SARS-CoV-2 virus 10-fold serial dilution: 10^1^ TCID_50_/well, 10^2^ TCID_50_/well, and 10^3^ TCID_50_/well. (**C-F)** Protection effect of EVs on LPS stimulation in lung fibroblast cells (LL24), bronchial epithelial cells (Beas-2B), microglial cells (BV2), and neuronal cells (NSCs). **(G)** Location of NF-κB p65 location after LPS treatment or LPS+EV treatment. **(H)** Expression of NF-κB p65 and the expression of phospho-p65 after LPS or LPS+EV treatment measured by western blot analysis. *, **, and *** indicate < 0.05, < 0.01, and < 0.001, respectively compared with the EV treatment group. †, ††, and ††† indicate p < 0.05, < 0.01, and < 0.001, respectively compared with the LPS treatment group. # indicates comparison between LPS and treated groups.

Viruses infecting the respiratory system can invade cells expressing the ACE2 receptor, resulting in cell damage. Because ACE2 receptors are expressed primarily in the liver, kidneys, male reproductive tissue, muscle, and the gastrointestinal tract (GI), these organs can be damaged by such viruses (**Suppl. Fig. 2A, B**). Although expression of ACE2 in the brain has not been studied in detail and the Brain Atlas database suggests that expression is very low (**Suppl. Fig. 2A, B**), clinical outcomes continue to suggest that the brain might be a target of SARS-CoV-2. We found that Neuronal stem cells (NSCs) strongly express ACE2 receptors (**Suppl. Fig. 2C**) at levels comparable with known virus target organs, such as LL24, indicating that the brain is a target organ for SARS-CoV-2. To determine whether EVs regulate pro-inflammatory cytokine release in response to SARS-CoV-2, we examined their indirect antiviral effects on these various cell types such as LL24, Beas-2B, and NSCs including brain immune cells (microglia cells; BV2), expected to be targeted by SARS-CoV-2. EVs labeled with PKH26 were observed after 24 h. Based on the results, the next experiments were conducted using an EV treatment time of 24 h (**Suppl. Fig. 2D**). Cells treated with EVs significantly reduced inflammation stimulated by Lipopolysaccharides (LPS). Expression of IL-1β and IL-6 in LL24 and Beas-2B cells (**Fig. 2C and D**, *Ps* < 0.05), of IL-1β, IL-6, and TNF-α in BV2 cells (**Fig. 2E**, *Ps* < 0.01), and IL-1β and IL-6 in NSCs (**Fig. 2F**, *Ps* < 0.05) fell significantly, whereas expression of IL-1β, IL-6, and TNF-α in cells not pre-treated with EVs increased after exposure to LPS. These results suggest that EV treatment may prevent cytokine storms caused by SARS-CoV-2 infection. NF-κB is a major transcription factor stimulated by LPS. Under normal conditions, NF-κB is located within the cytoplasm. Upon activation, the IKK complex phosphorylates IκBα, leading to translocation of NF-κB to the nucleus. Although NF-κB is an important regulator of immune responses, we found that expression of mRNA encoding NF-κB was not affected by EVs after LPS stimulation. Therefore, we examined the translocation of NF-κB protein. NF-κB was translocated into nucleus in LPS treated Beas-2B cells, but was sequestered in the cytoplasm after EVs treatment **(Fig. 2G)**. In addition, expression of NF-κB phospho-p65 in Beas-2B cells increased upon treatment with LPS and decreased after treatment with LPS and EVs; however, total p65 expression was unchanged (**Fig. 2H**). These results suggest that EVs regulate the inflammatory responses by blocking the NF-κB signaling pathway and p65 translocation.

### miRNAs in EVs directly bind to 3’ UTR of SARS-CoV-2

To find the mechanism of EV’s antiviral effect, we first analyzed miRNAs within EVs. Based on the fact that the viral genome can be targeted by human miRNAs, we hypothesized that miRNAs within EVs directly interact with the SARS-CoV-2 genome. EVs obtained from eight MSCs under various cell culture methods and from six placenta derivatives were assessed by small RNA sequencing. Quality control of EV sequencing results was performed using the qrqc R package. The distribution of small RNAs (expressed as a percentage) was obtained using sRNAtoolbox. Unassigned reads were most common, followed by miRNAs and tRNAs, indicating that miRNAs are the major cargoes in EVs (**Suppl. Fig. 3A-F**).

**Fig. 3.**
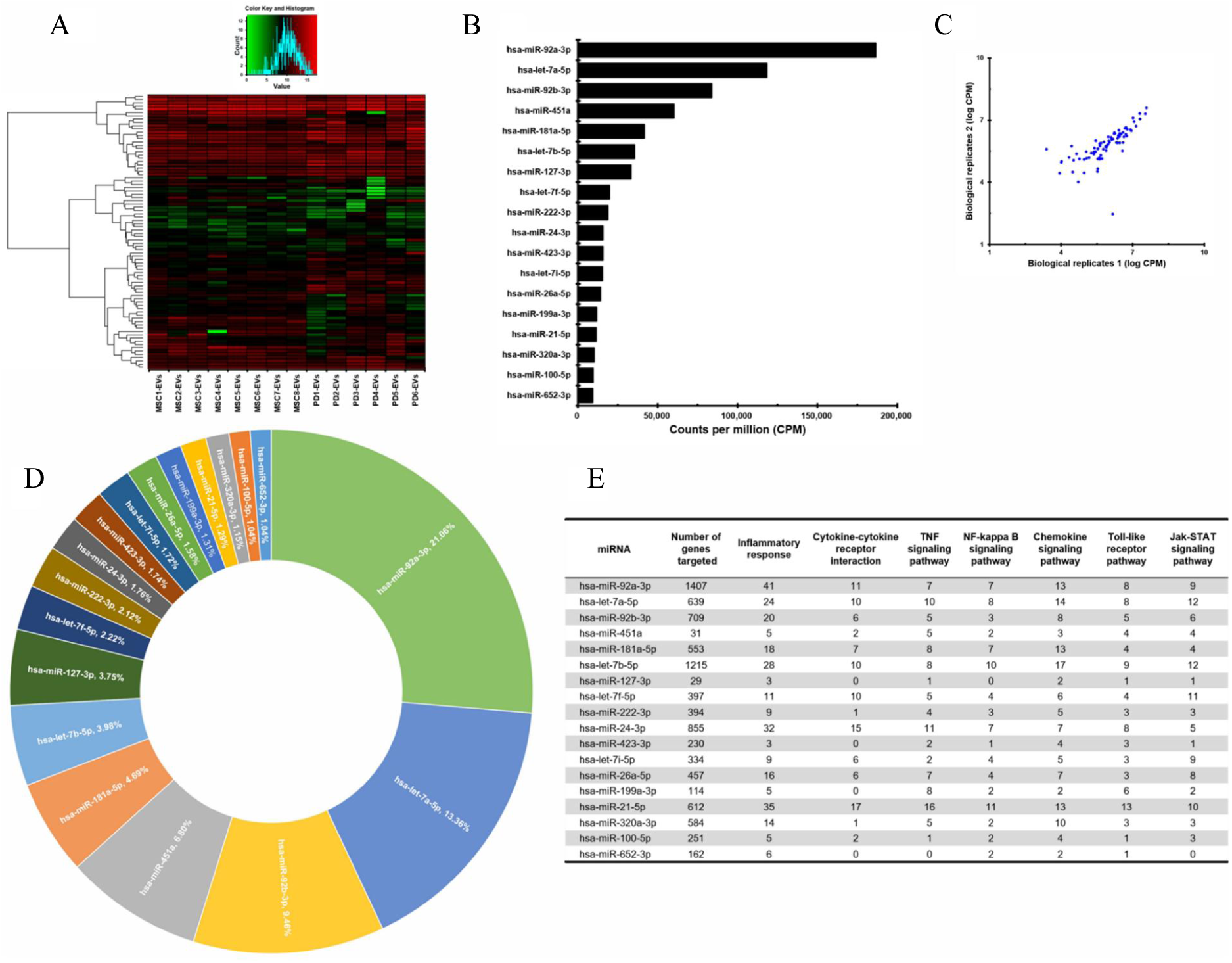
Profiles of miRNAs of pMSC-EVs and placenta EVs. **(A)** The miRNA profiles of eight MSCs and six placental derivatives EVs (PD-EVs). **(B)** pMSC-EV miRNAs were ranked (highest to lowest) in terms of counts per million (CPM). **(C)** A strong correlation between biological replicates. **(D)** The percentage of top 18 miRNAs accounted for all EV miRNAs. **(E)** 5380 genes that are predicted to be targeted by 18 miRNAs in a validated miRNA target database (the mirTarBase) and a breakdown of over-represented pathways and biological processes is shown.

The profiles of miRNA in EVs obtained from eight placenta MSCs and six placental derivatives EVs (PD-EVs) were very similar (**Fig. 3A**). MSC-EVs miRNAs were profiled using small RNA sequencing and ranked (highest to lowest) in terms of counts per million (CPM) (**Fig. 3B**). Read counts between biological replicates showed a strong correlation (**Fig. 3C**). The top 18 miRNAs accounted for 80.1% of all EV miRNAs (**Fig. 3D**). The remaining miRNAs were excluded from further analysis because they were present at very low read counts and comprised a very small percentage of the total reads (0.02–0.96%), and consequently were deemed unlikely to have significant biological effect relative to the more abundant miRNAs. The number of genes targeted by the top 18 miRNAs and the number of targeted genes related to the inflammatory response are listed in Figure 3E. In addition, the number of inflammatory response genes targeted was further assessed and the results of enriched pathway analysis were also shown in Figure 3E. The majority of the genes contributed substantially to specific inflammatory responses related to cytokine-cytokine receptor interactions, TNF, NF-KB, chemokines, Toll-like receptors, and the Jak-STAT signaling pathway, indicating that their key function is immunomodulation (**Fig. 3E**). The miRNAs affecting the most targets related to inflammatory responses were miR-92a-3p, miR-21-5p, miR-24-3p, miR-let-7b-5p, miR-let7a-5p, and miR-181a-5p. Moreover, the expression levels of 84 common miRNAs present in EVs derived from eight MSCs cultured under various conditions and six placental derivatives were very consistent across samples (**Suppl. Table 1**).

**Table 1.**
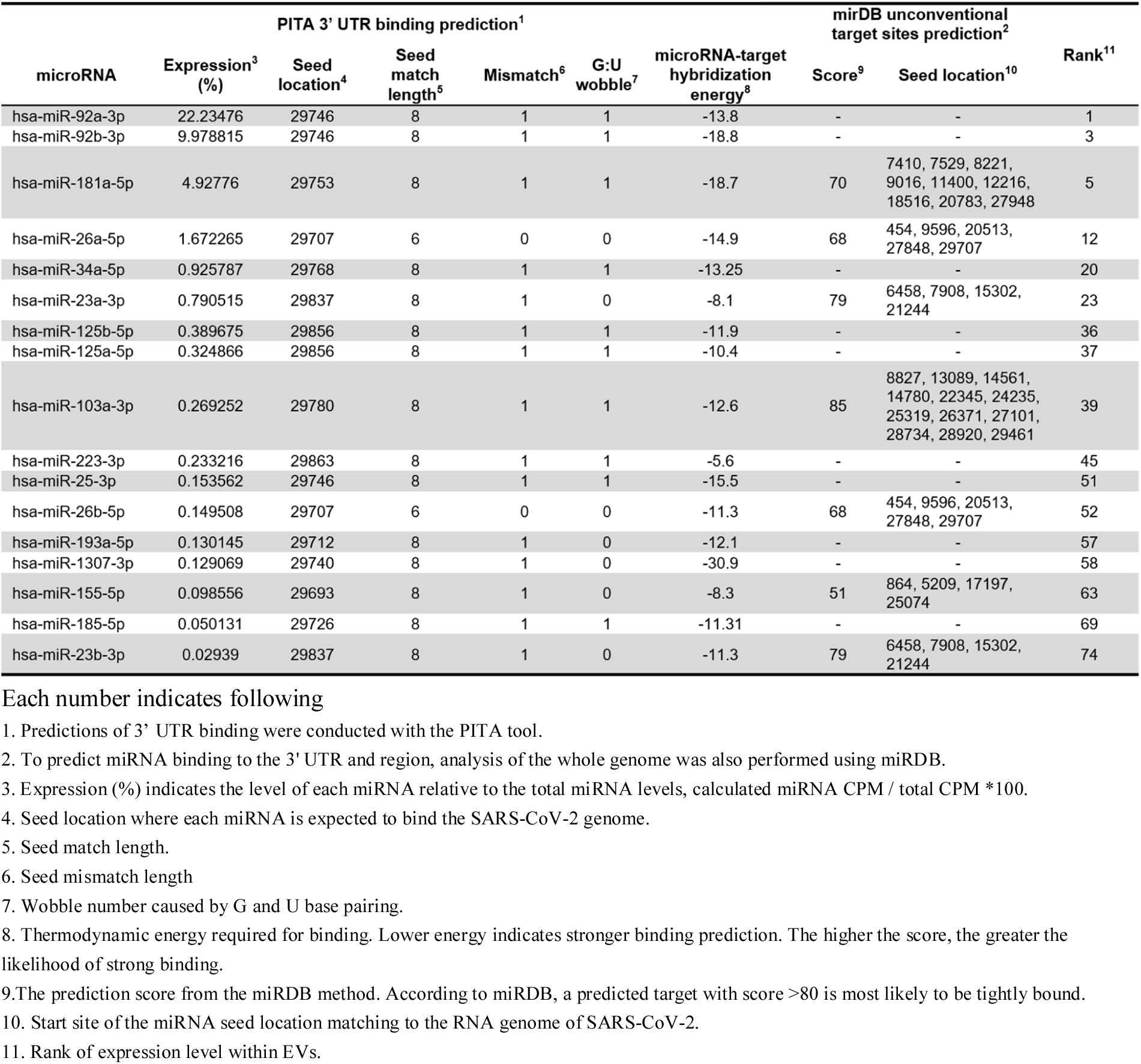
miRNAs predicted to bind the SARS-CoV-2 RNA genome.

| microRNA | Expression<sup>3</sup><br>(%) | PITA 3' UTR binding prediction<sup>1</sup> |  |  |  | microRNA-target<br>hybridization<br>energy<sup>8</sup> | mirDB unconventional<br>target sites prediction<sup>2</sup> |  | Rank<sup>11</sup> |
| --- | --- | --- | --- | --- | --- | --- | --- | --- | --- |
|  |  | Seed<br>location<sup>4</sup> | Seed<br>match<br>length<sup>5</sup> | Mismatch<sup>6</sup> | G:U<br>wobble<sup>7</sup> |  | Score<sup>9</sup> | Seed location<sup>10</sup> |  |
| hsa-miR-92a-3p | 22.23476 | 29746 | 8 | 1 | 1 | -13.8 | - | - | 1 |
| hsa-miR-92b-3p | 9.978815 | 29746 | 8 | 1 | 1 | -18.8 | - | - | 3 |
| hsa-miR-181a-5p | 4.92776 | 29753 | 8 | 1 | 1 | -18.7 | 70 | 7410, 7529, 8221,<br>9016, 11400, 12216,<br>18516, 20783, 27948 | 5 |
| hsa-miR-26a-5p | 1.672265 | 29707 | 6 | 0 | 0 | -14.9 | 68 | 454, 9596, 20513,<br>27848, 29707 | 12 |
| hsa-miR-34a-5p | 0.925787 | 29768 | 8 | 1 | 1 | -13.25 | - | - | 20 |
| hsa-miR-23a-3p | 0.790515 | 29837 | 8 | 1 | 0 | -8.1 | 79 | 6458, 7908, 15302,<br>21244 | 23 |
| hsa-miR-125b-5p | 0.389675 | 29856 | 8 | 1 | 1 | -11.9 | - | - | 36 |
| hsa-miR-125a-5p | 0.324866 | 29856 | 8 | 1 | 1 | -10.4 | - | - | 37 |
| hsa-miR-103a-3p | 0.269252 | 29780 | 8 | 1 | 1 | -12.6 | 85 | 8827, 13089, 14561,<br>14780, 22345, 24235,<br>25319, 26371, 27101,<br>28734, 28920, 29461 | 39 |
| hsa-miR-223-3p | 0.233216 | 29863 | 8 | 1 | 1 | -5.6 | - | - | 45 |
| hsa-miR-25-3p | 0.153562 | 29746 | 8 | 1 | 1 | -15.5 | - | - | 51 |
| hsa-miR-26b-5p | 0.149508 | 29707 | 6 | 0 | 0 | -11.3 | 68 | 454, 9596, 20513,<br>27848, 29707 | 52 |
| hsa-miR-193a-5p | 0.130145 | 29712 | 8 | 1 | 0 | -12.1 | - | - | 57 |
| hsa-miR-1307-3p | 0.129069 | 29740 | 8 | 1 | 0 | -30.9 | - | - | 58 |
| hsa-miR-155-5p | 0.098556 | 29693 | 8 | 1 | 0 | -8.3 | 51 | 864, 5209, 17197,<br>25074 | 63 |
| hsa-miR-185-5p | 0.050131 | 29726 | 8 | 1 | 1 | -11.31 | - | - | 69 |
| hsa-miR-23b-3p | 0.02939 | 29837 | 8 | 1 | 0 | -11.3 | 79 | 6458, 7908, 15302,<br>21244 | 74 |

Each number indicates following
1. Predictions of 3' UTR binding were conducted with the PITA tool. 2. To predict miRNA binding to the 3' UTR and region, analysis of the whole genome was also performed using miRDB. 3. Expression (%) indicates the level of each miRNA relative to the total miRNA levels, calculated miRNA CPM / total CPM \*100. 4. Seed location where each miRNA is expected to bind the SARS-CoV-2 genome. 5. Seed match length. 6. Seed mismatch length 7. Wobble number caused by G and U base pairing. 8. Thermodynamic energy required for binding. Lower energy indicates stronger binding prediction. The higher the score, the greater the likelihood of strong binding. 9. The prediction score from the miRDB method. According to miRDB, a predicted target with score >80 is most likely to be tightly bound. 10. Start site of the miRNA seed location matching to the RNA genome of SARS-CoV-2. 11. Rank of expression level within EVs.

To explore the possibility that miRNAs within EVs interact with the SARS-CoV-2 3’ UTR, we investigated whether the SARS-CoV-2 3’ UTR contains potential miRNA binding sites. We aligned the 3’ UTR sequences of SARS-CoV-2 isolates (**Fig. 4 A**) and analyzed the alignment using PITA software. Table 1 shows information about binding sites in the SARS-CoV-2 genome and the results of binding prediction analyses. PITA software identified 18 miRNAs predicted to target the 3 UTR of SARS-CoV-2 (**Table 1**) and 28 miRNAs with sequences expected to bind to whole genome sites within the SARS-CoV-2 (**Suppl. Table 2**). Five miRNAs: miR-92a-3p, miR-26a-5p, miR-23a-3p, miR-103a-3p, and miR-181a-5p out of 17 miRNAs were selected based on the thermodynamic energy score and high scores in two prediction tools: PITA and miRDB. Each of the candidate sites was assigned a logistic probability as a measure of confidence. We obtained the total free energy of each miRNA, based upon that the lower the binding thermodynamic energy (kcal/mol), the stronger the binding. For example, the binding energy of miR-181a-5p for the 3’ UTR was −18.7 kcal/mol, suggesting that binding of miRNA to the 3’ UTR binding would proceed spontaneously. PITA and miRDB predicted potential binding sites for miR-92a-3p, miR-26a-5p, miR-23a-3p, miR-103a-3p, and miR-181a-5p in the SARS-CoV-2 3’ UTR (**Fig. 4B**). Next, we performed qPCR analysis to determine whether these five miRNAs were present in EVs (**Fig. 4C**). The results confirmed that EVs expressed these miRNAs. To demonstrate the five miRNAs directly bind the SARS-CoV-2 3’ UTR and suppress the RNA replication, a luciferase reporter assay was developed. A PCR fragment was cloned into a luciferase reporter plasmid between the luciferase ORF and the synthetic poly(A) sequence. ‘recombinant plasmid was named pGL3 SARS-CoV-2 3’-UTR_Luc (**Fig. 4D**). The recombinant plasmid was transfected into human neuroblastoma SK-N-BE(2)C cells, and luciferase activity was measured 48 h later. As shown in Figure 4E, relative luciferase activity in SK-N-BE(2)C cells transfected with the recombinant plasmids was significantly lower than that in cells transfected with the empty psi-control vector (*Ps* < 0.001). Specifically, relative luciferase activity was downregulated when 3’ UTR was cotransfected along with expression vectors containing each of the five miRNAs or a vector containing all five miRNAs. The results show a significant decrease in the luciferase activity mediated by the psi-59 UTR (**Fig. 4E**). These data clearly indicate that each of the five miRNAs was able to silence the SARS-CoV-2 3’ UTR. Taken together, the results, suggest that all five miRNAs bind directly to the 3’ UTR of SARS-CoV-2 and degrade the SARS-CoV-2 viral genome. Based on these results, the five miRNAs tested all directly bind to the 3’ UTR of SARS-CoV-2 and degrade the SARS-CoV-2 viral genome further suggesting that either each of miR-92a-3p, miR-26a-5p, miR-23a-3p, miR-103a-3p, and miR-181a-5p or five miRNAs represent potential therapeutic agents for SARS-CoV-2.

**Fig. 4.**
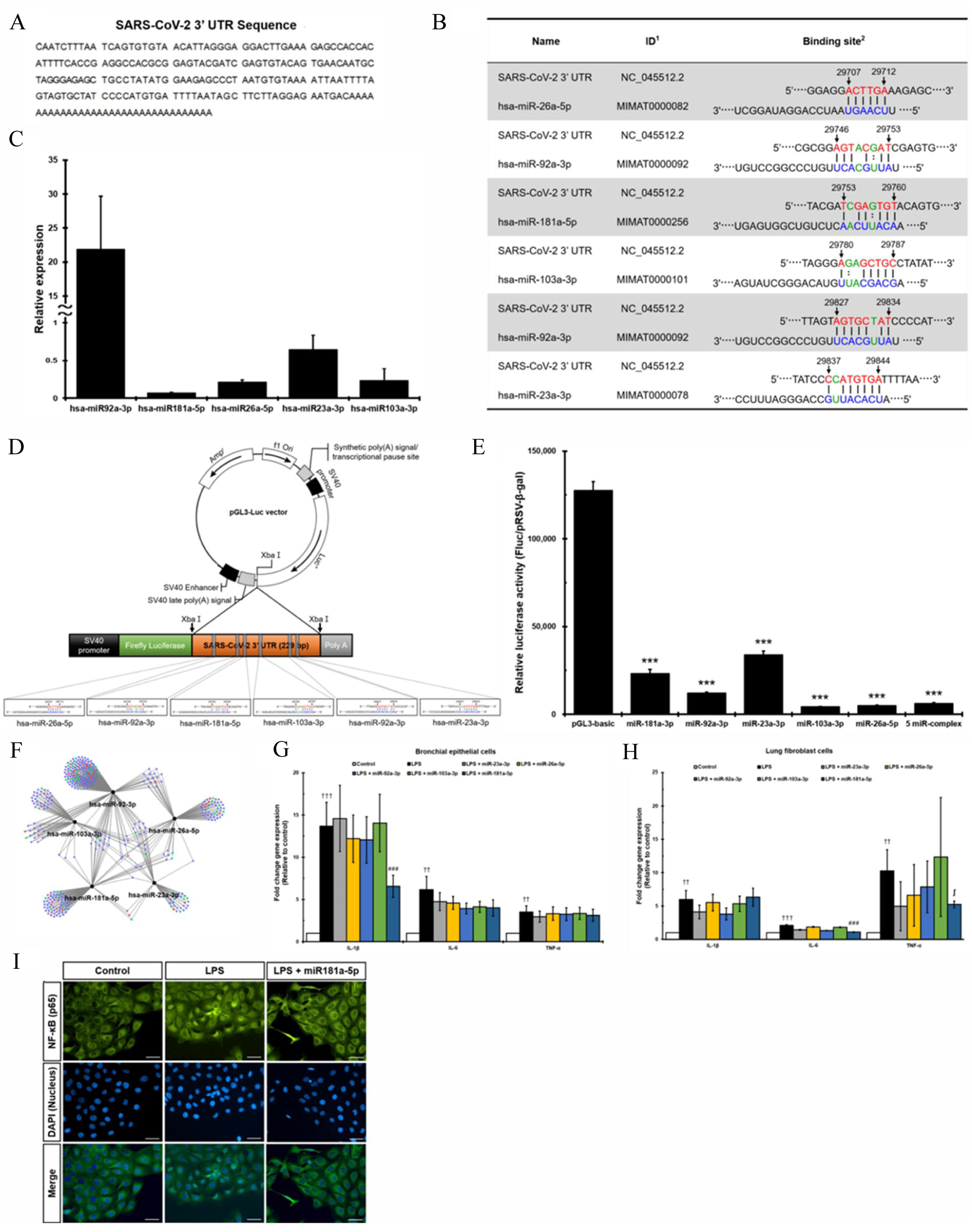
Direct viral effect of miRNAs in EVs on SARS-CoV-2. **(A)** The 3’ UTR sequence of SARS-CoV-2. The sequence was obtained from the NCBI database (accession NC_045512.2). **(B)** SARS-CoV-2 3’ UTR binding sites of five miRNAs. Each number indicates the followings. 1. Accession ID of NCBI or miRBase 2. The interaction diagrams of binding site in 3’ UTR of SARS-CoV-2. A line means perfect match, and a dot line means G:U wobble pairs. An empty space in seed regions means mismatch. **(C)** Confirmation of expression of five miRNAs measured by qPCR **(D)** Schematic representation of the luciferase constructs used for the reporter assays. Specifically, the pGL3 SARS-CoV-2 3′ UTR construct was transfected into human neuroblastoma SK-N-BE(2)C cells. Co-transfection of these cells with miR-92a-3p, miR-26a-5p, miR-23a-3p, miR-103a-3p, or miR-181a-5p, or all five miRNAs together, reduced luciferase activity. **(E)** The SARS-CoV-2 3’ UTR is a putative target of the five miRNAs identified in this study. The luciferase reporter assay revealed that miR-92a-3p, miR-26a-5p, miR-23a-3p, miR-103a-3p, and miR-181a-5p are potential regulators of SARS-CoV-2. **(F)** Enrichment analysis of biological processes targeted by the five miRNAs in EVs. The blue circle represents genes related to transcription, the green circle indicates immune regulation function, and the red circle shows genes with dual functions of transcription and immune regulation. **(G)** Expression of IL-1β, IL-6, and TNF-α. Bronchial epithelial cells were treated with each of miRNA, followed by LPS. **(H)** Expression of IL-1β, IL-6, and TNF-α. Lung fibroblast cells were treated with each miRNA, followed by LPS. **(I)** miR-181a-5p treatment dramatically reduces NF-κB translocation to nucleus. ∫, *, **, and *** indicate <0.06, < 0.05, < 0.01, and < 0.001, respectively compared with EV treatment group. †, ††, and ††† indicate < 0.05, < 0.01, and < 0.001, respectively compared with LPS treatment group. # indicates comparison between LPS and EV treated groups.

Enrichment analysis of biological processes targeted by the five miRNAs in EVs is shown in Figure 4F. Nodes represent individual genes associated with significantly enriched biological processes and the miRNAs that target them were connected by the edges. Genes related to transcription and immune regulation (including anti-inflammatory genes) are shown as blue and green circles, respectively, and genes with overlapping contributions to transcription and immune regulation are shown as red circles (**Fig. 4F**). Interestingly, all five miRNAs had dual roles in transcription and immune regulation, further confirming that they have the potential to both degrade SARS-CoV-2 via the direct binding with the 3’ UTR and regulation of the inflammatory environment created by viral infection. Based on the enrichment analysis of the five miRNA target genes in Figure 4F, the five miRNAs had the function of immune regulation. We further examined whether either each of the five miRNAs within the EVs regulate pro-inflammatory cytokine release in response to SARS-CoV-2. LL24 and Beas-2B were exposed to either each of five or all of miRNAs, followed by LPS treatment. Expression of pro-inflammatory cytokines was measured. Only miR-181a-5p exhibited significant reduction of IL-1β in Beas-2B (**Fig. 4G**, p < 0.0003); and IL-6 in LL-24 cells (**Fig. 4H**, p < 0.0005); the other four miRNAs did not reduce expression of pro-inflammatory factors. Interestingly, the increase in TNF-α induced by the inflammatory response was marginally significantly reduced by miR-181a-5p in Beas-2B (**Fig. 4G**, p < 0.01) and in LL-24 (**Fig. 4H**, p < 0.06). Furthermore, we found transfection of miR-181a-5p reduced NF-κB translocation to the nucleus markedly in Beas-2B cells (**Fig. 4I**). Next, we analyzed the processes and pathways targeted by the five miRNAs. The mirTarBase predicted that the five miRNAs targeted 2698 genes. GO term analysis using DAVID indicated that the functions of the targeted genes involved both positive and negative regulation of transcription from RNA polymerase II promoters, as well as other transcription-related functions (**Suppl Fig. 4A and B**). KEGG pathway analysis of the 2698 targets revealed significant enrichment of pathways related to aging (cell cycle, p53) and inflammation (PI3K, Wnt, TGF). We detected significant enrichment in pathways related to inflammatory cytokines such as TNF-α, cytokine-cytokine receptors, and chemokine receptors. Interestingly, RNA degradation, RNA transport, protein processing in the ER, and TGF-β were also enriched (**Suppl. Fig. 4C-F**). Some RNA viruses target the nuclear pore complex, thereby inhibiting RNA transport. The inhibition of RNA transport in the nuclear pore complex prevents influx of new mRNA into the cytosol, thereby reducing competition for the translation machinery and attenuating responses to viral infection [22]. By preventing suppression of RNA transport, which is a survival strategy for some viruses, the five miRNAs might prevent formation of an environment favorable for viral replication. The RNA degradation pathway mainly targets genes related to the CCR4 NOT complex, which is involved in RNA homeostasis and removal of unnecessary mRNA from cytoplasmic exosomes. Given that inflammation-specific mRNAs are not cleared from CNOT3 knockout mice [23], the five miRNAs may stimulate clearance by targeting those RNAs for degradation (**Suppl. Fig. 4D**). Upon infection with hepatitis C virus, ER stress markers are upregulated and proapoptotic markers are expressed [24]. The five miRNAs regulated expression of genes related to ER stress caused by virus infection (**Suppl. Fig. 4E**). Influenza A virus increases expression of TGF-β and promotes cell proliferation, thereby increasing the number of cells expressing integrin subunit α5 and fibronectin, to which bacteria adhere. In addition, the innate immune system suppresses expression of the TGF-beta receptor and the SMAD pathway via IRF3 during virus infection [25, 26]. By preventing suppression of this pathway, the five miRNAs normalized the local environment around virus-infected cells (**Suppl. Fig. 4F**). Of the 2698 targets, we selected 83 genes associated with the GO term ‘inflammatory response’, and then performed KEGG pathway analysis to further identify the pathways in which they are involved: these were interleukin production and regulation, cell chemotaxis, and response to external stress (**Suppl. Fig. 4A and B**). Moreover, the 83 genes were implicated in signal pathways related to cytokine production (e.g., the NF-κB pathway, the JNK pathway, and the toll-like receptor pathway) (**Suppl. Fig. 5A-G**). In addition, they target important inflammation-inducing factors such as IL-6, IL-8, and COX2. These data indicate that the five miRNAs regulate cytokine-mediated inflammatory responses and inflammatory cell activation, as well as pathways related to cytokine secretion, thereby dampening the inflammatory environment. Taken together, the above data suggest that the five miRNAs regulate the inflammatory environment and degrade viral genomic RNA through direct binding to the 3’ UTR of SARS-CoV-2.

**Fig. 5.**
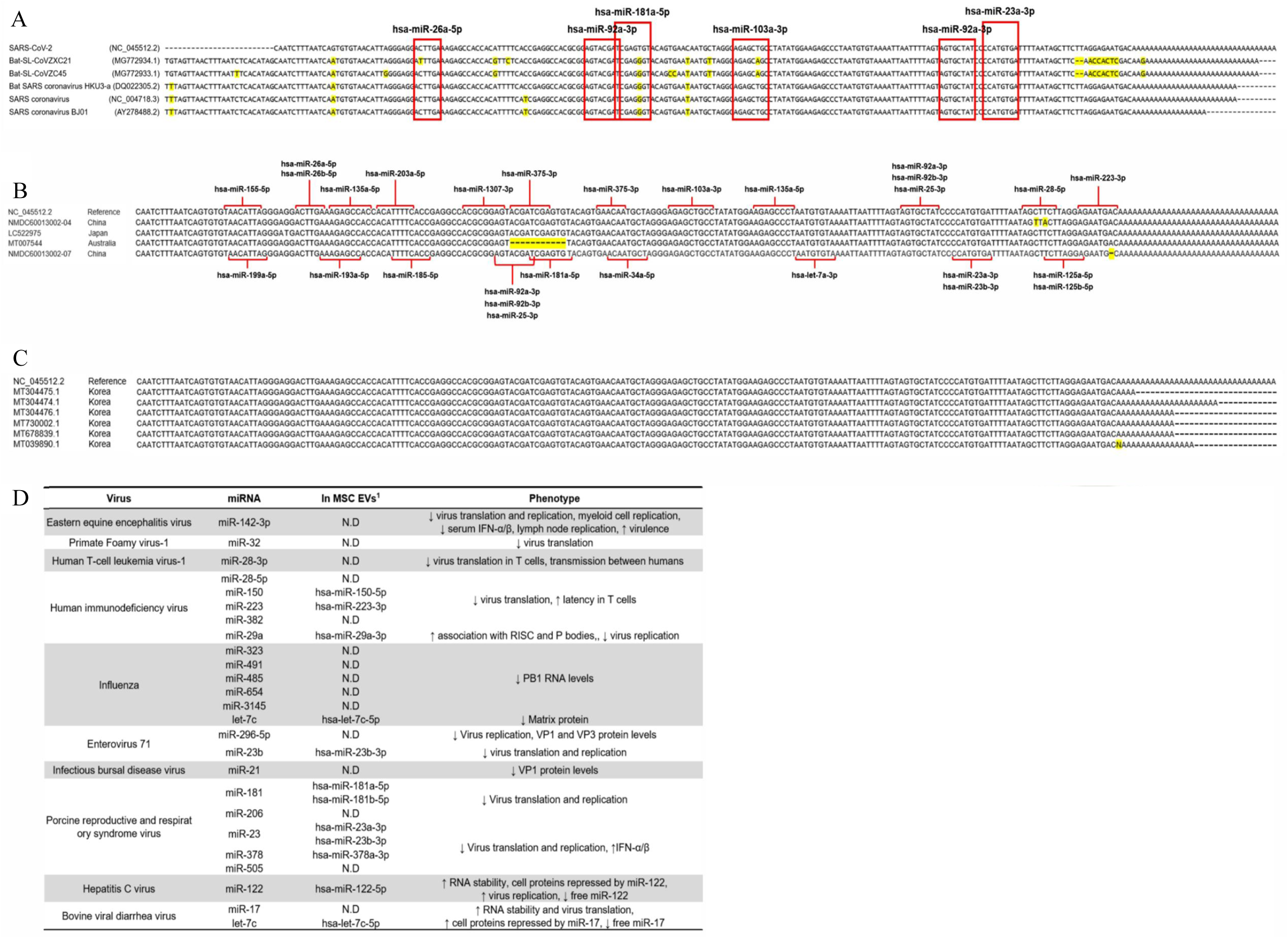
Multi-binding miRNAs will be able to target new mutant SARS-CoV-2 variants. **(A)** 3’UTR sequence of SARS-CoV-2 of selected five corona viruses. The red boxes indicate miRNA binding sites to the conserved regions within the 3’UTRs of five corona viruses. The label on the left panel showed virus names and GenBank accession. Mutated nucleotides are indicated in yellow. **(B)** Each 3’UTR sequence was aligned to 3 UTR sequence of NC_045512.2 using the MUSCLE tool. The label on the left panel showed GenBank accession and location where the virus was found. Red lines indicated interaction between 3 UTR sequence and miRNA. Mutated nucleotides are indicated in yellow. **(C)** 3 UTR sequence of 6 complete genome obtained from SARS-CoV-2 isolated form Korean patients was aligned. 6 complete genome of SARS-CoV-2 isolated form Korean patients were submitted to GenBank from 2020-February until 2020-July. Mutated nucleotides are indicated in yellow. **(D)** Modified table obtained from published reports (Trobaugh, Derek W., and William B. 2017). The table shows that known interaction between miRNA and viral RNA and there are several miRNAs existing in pMSC-EVs. 1 indicates the detected miRNAs in pMSC-EVs.

### 3’ UTRs of corona virus are highly conserved

Next, since the 3’ UTR sequence is conserved among corona viruses, and considering the hypothesis that a functional miRNA target site would be conserved across all corona viruses, we selected 3’ UTR sequences from five random coronaviruses and compared them using the MUSLCE tool [27]. The sequence of SARS-CoV-2 3’ UTR of SARS-CoV-2 was aligned against the other coronaviruses. The red boxes indicate miRNA binding sites and sites predicted to bind the five miRNAs were also highly conserved (**Fig. 5A**).

To confirm that the SARS-CoV-2 genomic mutation also occurs in the miRNA binding site of the 3’ UTR, the site of the 3’ UTR sequence in the recently generated mutant SARS-COV-2 was obtained from data reported in a recent paper [28]. According to, mutation of the 3’ UTR is found rarely in SARS-CoV-2 samples collected from China and Japan, whereas samples from Australia showed deletion of the 3’ UTR (**Fig. 5B**). Specifically, among the mutations, a total of 5 cases with 3’ UTR mutation were detected: 29705G>T, 29854C>T, 29856T>A, 29869del and 29749-29759del with the most common type mutation was single nucleotide mutation. Most sites within the SARS-CoV-2 3’ UTR that bind miRNAs are not mutated, whereas the binding sites of hsa-miR-181a-5p, hsa-miR-92a-3p, hsa-miR-92b-3p, and hsa-miR-25-3p were mutated in the SARS-CoV-2 samples from Australia. However, in the case of hsa-miR-92a-3p, two binding sites are predicted in the 3’ UTR; one of these binding sites is known to harbor none of the mutations found in any of the variants in China, Japan, and Australia. These results suggest that most miRNAs are rarely affected by known 3’ UTR mutations (**Fig. 5B**). Next, to investigate the SARS-CoV-2 complete genome from Korean patients, we used NCBI GenBank. Mutation of 3’ UTR was not detected in six complete genome of SARS-CoV-2 isolated form Korean patients which were recently submitted to GenBank from 2020-February until 2020-July. (**Fig. 5C**).

In summary, considering the high miRNA binding rate, and that the 3’ UTR of various coronaviruses rarely harbor mutations, the five selected miRNAs have the potential to suppress all coronaviruses, including those arising via mutation. Furthermore, several miRNAs in MSC-EVs interact with other viruses. Specifically, miR-150-5p, miR-223-3p, and miR-29a-3p suppress HIV translation and prolong its latency in T cells. In addition, Let-7c-5p reduces expression of matrix protein, which is critical for influenza virus, and miR-23b-3p blocks translation of Enterovirus 71. miR-181a-5p, miR-181b-5p, miR-23a-3p, miR-23b-3p, and miR-378a-3p degrade porcine reproductive and respiratory syndrome virus, whereas miR-122-5p and let-7c-5p suppress hepatitis C virus and bovine viral diarrhea virus, respectively (**Fig. 5D**). Consequently, the presence of miRNAs in MSC-EVs that are capable of attacking various RNA-based viruses suggests that MSC-EVs can be used to treat and/or prevent virus infection.

## Discussion

As yet, there is no vaccine or treatment for SARS-CoV-2. Consequently, symptomatic and maintenance therapies are the standard of care. While an effective and safe treatment for patients with SARS-CoV-2 infection is required urgently, SARS-CoV-2 is an RNA virus and as such is likely to undergo frequent mutation. Therefore, vaccines or treatments must also be effective against newly arising mutant viruses. Here, we demonstrate that miRNAs in EVs suppress SARS-CoV-2 by binding directly to the 3’ UTR, resulting in translational repression (**Fig. 3**). EVs prevent cytokine storms by normalizing inflammatory responses. In addition, each individual (or all five together) miRNA is predicted to interact directly with the 3’ UTR of SARS-CoV-2, which was confirmed by significant inhibition of a luciferase reporter construct derived from the 3’ UTR (**Fig. 4**). Moreover, EVs or miRNAs modulated host factors to normalize immunopathogenesis. More specifically, 84 miRNAs present in EVs exerted potent indirect effects by targeting mRNAs that stimulate the immune response. Binding of target pro-inflammatory mRNAs, along with the anti-inflammatory factors contained in EVs, could normalize the immune environment, and prevent cytokine storms.

Recent studies of the antiviral effects of miRNAs reveal that miRNA in host cells interacts directly with viral RNA to prevent its replication [29]. In addition, even if miRNAs do not interact directly with viral RNA, they can still suppress expression of viral target receptors, thereby subverting virus cell tropism. Furthermore, these miRNAs can regulate the replication and translation machinery involved in virus replication and inhibit viral infection by controlling inflammation and apoptosis, both of which promote the spread of assembled virus particles [30]. In addition, because miRNAs have sequences complementary to their targets, efforts have been made to control only specific target mRNAs using miRNAs. Exogenous miRNA is stable *in vivo* and is able to regulate its target [29]. However, studies of drug delivery systems are actively being conducted because it is necessary to localize miRNAs to the target tissue, delivery it to cells, and avoid nuclease degradation. To address these issues, some groups have tried modifying miRNAs. However, when such modifications are made, they create other problems, including off-target effects, reduction in miRNA activity, and formation of toxic metabolites due to miRNA degradation [31]. Consequently, a miRNA (small RNA)-based treatment was not released until 2018, when the FDA approved a drug called Patisiran (ONPATTRO, NCT01960348). Currently efforts are underway to develop lipid nanoparticles that can be delivered to target cells following intravenous (IV) injection (hepatocytes) allowing the efficient transfer of miRNA [32]. As research on extracellular vesicles advances, the paracrine and regenerative effects of stem cells mediated via EVs have been demonstrated [33]. Because EVs are natural vesicles they are considered to be safer and more effective than artificial alternatives.

Therapeutic agents based on miRNAs derived from EVs have several advantages as therapeutic agents. First, a major problem with conventional treatments for RNA-based viruses is that they have potentially fatal side effects. Clinical trials conducted to date show that existing viral treatments have strong side effects [34], making them inappropriate for treatment of patients with mild cases; moreover, the side effects make it difficult to give antiviral drugs in doses high enough to yield the desired effect. In contrast, due to specific binding to the 3’ UTR or to the SARS-CoV-2 genome, miRNAs derived from EVs are unlikely to have significant side effects. Therefore, it is necessary to overcome the side effects of existing antiviral agents and to develop more virus-specific drugs. As mentioned above, a miRNA-based agent has already been approved by the FDA, and the interaction between miRNA and RNA viruses has been proven, which makes therefore, the possibility of treatment is high. Moreover, because mutations in the 3’ UTR of the virus are very rare, this approach has the potential to become a universal treatment for novel RNA viruses arising via mutation of the viral RNA. Our study demonstrates that the 3’ UTR sequence is highly conserved across the corona virus family (**Fig. 5A**), and that binding sites predicted to bind the five miRNAs are also highly conserved (**Fig. 5A and B**). These results give us confidence that the five selected miRNAs will be effective against novel viruses arising though mutation.

Third, in addition to the direct antiviral effects of miRNA, MSC-EVs can transport various cargoes that can regulate immunity and promote regeneration of damaged tissue.

MSCs have attracted attention as COVID-19 therapeutics because several lines of evidence show that they can significantly improve the pathological state of damaged lungs, including pneumonia; in addition, they have the ability to activate phagocytosis by macrophages, thereby preventing virus spread [35]. Furthermore, because MSC-EVs contain factors that can regenerate stem cells, they exhibit some of the biological properties of MSCs, and express surface receptors, signal transduction molecules, cell adhesion molecules, miRNAs, and antigens characteristics of MSCs [36, 37], they will have effects against pneumonia similar to those of MSCs [38].

Furthermore, MSCs regulate the function of immune cells and accordingly are considered potential treatments for autoimmune and inflammatory diseases. According to numerous experimental and clinical studies, most MSC-based immunomodulatory functions can be attributed to the immunomodulatory properties of the EVs that they secrete. MSC-EVs are rich in biologically active molecules such as mRNA, miRNA, cytokines, and immune modulators, which regulate the function, phenotype, and viability of immune cells [39]. MSC-EVs with immune regulatory activity are also used to study degenerative, chronic, and infectious diseases. For example, when pigs infected with the influenza virus were treated with MSC-EVs, virus outflow (as detected by nasal swab) decreased, virus replication in the lungs diminished, and pro-inflammatory cytokine production was reduced significantly [40]. Because host immune responses in severe COVID-19 patients are exacerbated by pro-inflammatory factors such as IL-1, IL-6, IL-8, and impaired type I interferon activity, which can trigger a potentially fatal cytokine storm, a hallmark of severe COVID-19 according to several reports, they provide a rationale for therapeutic approaches based on EVs. In addition, a recent study reported that a multi-system inflammatory syndrome in children infected with SARS-CoV-2 results in a life-threatening disease. Therefore, it is very critical to control the inflammatory syndrome before and after virus infection [41].

Indeed, our study demonstrates that EVs from MSCs and various placental derivatives successfully inhibit secretion of pro-inflammatory factors (**Fig. 2, 4, and suppl Fig. 4 and 5**). The genes targeted by each of the top miRNAs contribute substantially to specific inflammatory responses (**Fig. 3E**) related to cytokine-cytokine receptor interactions, TNF secretion, NF-KB activation, chemokine secretion, Toll-like receptor expression, and the Jak-STAT signaling pathway, indicating that their key function is immunomodulation (**Suppl Fig. 5**). Specifically, miR-let-7b-5b, miR-21-5p, miR-92a-3p, miR-24-3p, and miR-181a-5p bind the most targets related to inflammatory responses.

MSC-EVs have several other advantages over MSCs [39]. First, MSC-EVs are easier to manipulate and store [42]. Also, because MSC-EVs are cell-free and nano-sized, they are far less likely to trigger immunogenicity, tumorigenicity, or embolism. Because they are essentially liposomes, EVs are simple biological structures that are more stable *in vivo* than other particles [43]. Moreover, EVs invade host cells directly through membrane fusion, receptor-mediated phagocytosis, and other internalization mechanisms, which contribute to physiological and pathological processes such as activation of signaling pathways and immune responses [44]. Taken together, the observations reported herein suggest that MSC-EVs are an effective antiviral treatment.

More interestingly, some of miRNAs in EVs are possible therapeutic agents for other RNA viruses **(Fig. 5B)**. Miravirsen illustrated that miRNAs that interact directly with RNA viral genomes can be used to treat hepatitis C infection; indeed, the drug has successfully completed a phase II clinical trial. Moreover, because various miRNAs present in MSC-EVs can be used to target RNA genomes with complementary sites in their 3’ UTRs, they could be used to treat infections caused by other coronaviruses, hepatitis viruses, and HIV-based RNA viruses **(Fig. 5C**).

Although MSC-EVs are expected to be effective for antiviral therapy and immune enhancement, one major obstacle to their widespread use is that they are difficult to mass-produce [45]. To mass-produce EVs consistently, it will be necessary to establish reproducible culture conditions and scale up the system in an economically viable manner. Moreover, mass production of EVs is limited by the challenge of producing MSCs themselves. However, in this study, we showed that the miRNA profiles of MSC-EVs and placental derivatives EVs are very similar. Consequently, it should be possible to obtain more EVs than cells by isolating placental derivatives EVs. Alternatively, it might be possible to synthesize effective miRNAs and package them into EVs to increase treatment efficacy.

To summarize, we demonstrated that EVs and five major miRNAs significantly inhibit SARS-CoV-2 replication and exert anti-inflammatory activity *in vitro*. Moreover, EVs regulate the pro-inflammatory environment induced by viral infection and suppress replication of SARS-CoV-2. In addition to the antiviral effect of EVs, EVs and miRNAs prevented spread of the virus when administered prior to a pro-inflammatory agent (LPS).

## Materials and Methods

### Isolation of human brain neuronal stem cells and placental MSCs

Neuronal stem cells were generated from the central nervous system tissue of spontaneously fetus from ectopic pregnancy at during the gestational week (GW) 8 with mother’s consent (IRB 2009-06-074). Isolation method is described in Moon et al [46]. Normal human placenta (≥37 GW) showing no evidence of medical, obstetrical, or surgical complications were obtained. All donors provided written, informed consent. Sample collection and use for research purposes were approved by the Institutional Review Board (IRB, 2015-08-130-016) of Bundang CHA General Hospital (Seongnam, Korea).

### Isolation of EVs

EVs were first isolated by centrifugation at 4,000 × g for 30 min using in an 10K MCWO spin column (Merk Millipore, Burlington, MA, USA) and then filtered through a qEV original size exclusion column (Izon Science Ltd, Burnside, Christchurch, New Zealand). The qEV original size exclusion column was removed from storage at 4°C and the 20% ethanol storage solution was allowed to run through the column followed by 20 mL particle-free PBS. Samples were diluted to 500 µL with particle-free PBS and then overlaid on the qEV size exclusion column, followed by elution with particle-free PBS. The flow was collected in 500 µL fractions, and EV-rich fractions 7 and 10 were combined for analysis. The combined fractions were centrifuged at 1,000 × g for 1 min in an Ultrafree 0.22 μm centrifugal filter device (Merk Millipore).

### Fluorescence imaging of EVs

EVs were biotinylated using EZ-Link Sulfo-NHS-LC LC-Biotin (Thermo Fisher Scientific, Waltham, MA, USA). Biotinylated EVs were loaded onto Zeba spin desalting columns (7 K MWCO; Thermo Fisher Scientific) to remove the remaining free biotin. For staining, 20 μL of biotinylated EVs were placed in a circle drawn on a streptavidin-coated glass slide (Arrayit Corporation, Sunnyvale, CA, USA). After 30 min, EVs were fixed with BD Fix Perm (BD Biosciences, San Jose, CA, USA) and blocked with 0.2% BSA-PBS. Immunofluorescence staining was performed using anti-human CD 63 (1:100; SC-5275, Santa Cruz Biotechnology, Dallas, TX, USA) for 2 h at room temperature, followed by incubation with Alexa Fluor 488-conjugated secondary antibodies (1:500; Invitrogen, Carlsbad, CA, USA) for 1 h at room temperature.

### TEM imaging

Five-microliter aliquots of diluted sample containing 0.5 µg protein (as determined by BCA assay) were dropped onto hydrophilic grids. After a few minutes, the grids were washed with distilled water and contrasted for 20 sec with 2% uranyl acetate. Images were acquired on a JEM-1010 microscope (JEOL, Seoul, Korea) operating at 80 kV.

### Nanoparticle tracking analysis

EVs were diluted in PBS (final volume, 1 mL) and examined under a ZetaView Nanoparticle Tracking Video Microscope (Particle Metrix, Inning, Germany). The software manufacturer’s default settings for EVs or nanoparticles were selected. For each measurement, three cycles were performed by scanning 11 cell positions and capturing 60 frames per position (video setting: high). The following settings were used: focus, autofocus; camera sensitivity for all samples, 80.0; shutter, 100; scattering intensity, 1.2; cell temperature, 23°C. After capture, the videos were analyzed using the built-in ZetaView Software 8.02.31 with the following parameters: maximum particle size, 1000; minimum particle size, 5.

### Western blotting

EVs were lysed in 10× RIPA buffer containing a protease cocktail tablet (Roche, Basel, Switzerland) and phosphatase inhibitors II and III (Sigma-Aldrich). Protein concentrations were measured using the BCA assay (Thermo Fisher Scientific). A 20 μg aliquot of each sample was subjected to 10% SDS-PAGE, followed by western blotting. Blots were blocked for 1 h in 10% skim milk/TBS-T. Antibodies specific for the following proteins were obtained from the indicated suppliers: CD81 (1:1,000, Santa Cruz Biotechnology, Dallas, TX, USA), CD9 (1:1,000, Santa Cruz Biotechnology), CD63 (1:1,000, Santa Cruz Biotechnology) TSG 101 (1:500, Abcam, Cambridge, UK), Vinculin (1:2,000, Abcam). Cells were lysed in 1× RIPA buffer containing a protease cocktail tablet and phosphatase inhibitors II and III. Next, 10 μg of protein was loaded onto 10% SDS-PAGE gels, transferred to a PVDF membrane (Merck Millipore), and analyzed by western blotting with antibodies specific for p65 (1:1,000, Cell Signaling Technology, Danvers, MA USA), phosphor-p65 (1:1,000, Cell Signaling Technology), and GAPDH (1:10,000, Santa Cruz Biotechnology). Primary antibodies were diluted in TBS-T and incubated with blots overnight at 4°C, followed by incubation with anti-rabbit or anti-mouse horseradish peroxidase (HRP) conjugated secondary antibodies (Jackson immunoresearch, West Grove, PA, USA) for 1 h. Immunoreactivity was detected using enhanced chemiluminescent HRP substrate (Merck Millipore).

### Cells and viruses

The African green monkey kidney epithelial cell line (Vero) was used as an appropriate cell line for SARS-CoV-2 propagation. The cells were cultured in Dulbecco’s Modified Eagle’s Medium (DMEM, Gibco) supplemented with 10% fetal bovine serum (FBS, Gibco). Cultured Vero cells were incubated at 37°C/5% CO_2_ in a humidified chamber. The antiviral activity assay was carried out using the NMC-nCoV02 strain of SARS-CoV-2. The virus was propagated in Vero cells and the titer of propagated viral stock was expressed as the tissue culture infective dose 50 (TCID_50_). All experiments using the virus were performed in a Biosafety level-3 (BLS-3) laboratory located in College of Medicine and Medical Research Institute, Chungbuk National University, Cheongju 28644, Republic of Korea. Cell lines and virus were provided by Chungbuk University.

### Cytotoxicity assay

Cytotoxicity values of the EVs on Vero Cells were determined using the Methyl Thiazolyl Tetrazolium (MTT) assay. Briefly, 1 × 10^4^ of Vero cells in 96-well cell culture plates were treated with different concentrations (2-fold serial dilutions) of each EV in triplicate. Treated cells were incubated for 24 h at 37°C, followed by addition of 5 mg/mL of MTT solution per well. The microplate was incubated at 37°C for 4 h. Then, 100 μL of DMSO solution was added to each well. Cell viability was calculated after measuring the absorbance (at 500-600 nm) in a micro plate reader.

### TCID_50_ assay

The 1 × 10^4^ of Vero cells were seeded into 96-well plate and incubated overnight in 37°C/5% CO_2_ humidified incubator. Then, the SARS-CoV-2 were serially diluted 10-fold and 0.1 ml of each dilution was added to triplicate wells of 96-well plates containing confluent Vero cells (final volume 0.2 mL/well). After 3 days, cells were washed by PBS and fixed by 10% formaldehyde solution for 5 minutes and stained by 1% crystal violet. The wells were inspected for the presence of virus, as judged by the appearance of CPE (Cytopathic Effect). The virus endpoint titration (dilution required to infect 50% of the wells) was expressed as TCID_50_/mL. CPE (%) was calculated, along with the virus titer (number of positive wells/total number of wells × 100).

### Antiviral activity assay

The cytotoxicity inhibition assay was used to the antiviral activity of each EVs. Briefly, Vero cells were grown in a 96-well plate (1 × 10^4^ cells/well) at 37°C/5% CO_2_ until 80% confluent. Subsequently, the culture medium was removed from each well and 10^1^–10^2^ TCID_50_ of COVID virus (50 μL of suspension) and different concentrations (2-fold serial dilutions) of EVs (50 μL) were added to each well. For the virus control, 10^1^–10^3^ TCID_50_ of virus plus the highest concentration of DMSO were added to six wells. Also, six wells were treated with DMSO alone (negative control). The plates were incubated at 37°C/5% CO_2_ and CPE monitored for up to 3 days post-infection.

### Cell culture

The Human Lung fibroblast cell line LL24, Human Bronchial epithelial cell line Beas-2B, Mouse microglial cell line BV2 were obtained from ATCC (Manassas, VA, USA). LL24 cells were grown at 37°C in a humidified atmosphere of 5% CO_2_ in RPMI-1640 medium (Gibco), supplemented with 10% FBS (Gibco) and 1% penicillin/streptomycin (P/S, Gibco). Passage was performed every 3 days. Beas-2B cells were cultured at 37°C in a humidified atmosphere of 5% in serum-free 1× defined keratinocyte SFM Gibco) supplemented with 5 µg/mL gentamicin (Gibco). The medium was replaced every 2–3 days and cells were passaged every 4–5 days. BV2 were cultured at 37°C/5% CO_2_ in growth medium (Dulbecco’s modified Eagle’s medium (D-MEM, Gibco), supplemented with 10% FBS and 1% P/S. BV2 were passaged every 2-3 days. NSCs were isolated from human fetal tissue at GW 10, 12, and 14 with the mother’s consent [46]. Neurons were cultured at 37°C, 5% CO_2_, and 3% O_2_ in DMEM/F12 (Gibco) with B27 supplement (Gibco), 50 µg/mL gentamicin (Gibco), human bFGF, human EGF 20 ng/mL (PeproTech, Rocky Hill, NJ, USA), tocopherol, and tocopherol acetate (1 µg/mL; Sigma-Aldrich).

### PKH26 labeling of EVs

EVs were labeled with the PKH26 Red Fluorescent Cell Linker Kit for General Cell Membranes (Sigma-Aldrich). Briefly, EV pellets were resuspended at 200 µg/mL in diluent C and then mixed 1:1 with dye solution (2 µL PKH26 ethanolic dye and 500 µL diluent C) for 5 min. Next, an equal volume of 1% bovine serum albumin (BSA, Merk Millipore) was added to bind excess dye and the resultant solution was passed through 7 K MWCO Zeba Spin Desalting Columns (Thermo Fisher Scientific) to remove excess dye. Samples were stored at −80°C.

### LPS and EV treatment

For experiments, cells were seeded at equal densities into wells of 6-well plates (Nunc, Roskilde, Denmark). The cells were pre-treated with EVs for 24 h and then stimulated with LPS (LL24: 2 μg/mL, Beas-2B: 4 μg/mL, BV2 and NSCs: 1 μg/mL) for either 6 h or 1h.

### Immunocytochemistry (ICC)

Beas-2B cells were cultured and fixed in 4 % PFA for 30 min. Cells were washed three times with PBS and blocked with 1 % BSA in PBS for an additional 20 min at room temperature. Cells were then incubated with anti-NF-κB p65 subunit (1:50, Santa Cruz Biotechnology) for 24 h at 4°C. Following washing with PBS three times to remove excess primary antibody, the cells were further incubated with Alexa Fluor 488-conjugated goat anti-mouse IgG (1:2,000; Invitrogen) for 1 h at room temperature, washed with PBS, and stained with Gold Antifade reagent containing DAPI (Invitrogen; Thermo Fisher Scientific.) for 5 min prior to the position and quantification of ProLong nuclei. Subsequently, the slides were coverslipped and visualized using fluorescence microscope (Nikon, Minato City, Tokyo, Japan).

### RNA extraction and quantitative PCR (qPCR)

Total RNA for qPCR was extracted from LL24, Beas-2B, BV2 and NSCs. mRNA was extracted from EVs using the TRIzol reagent (Invitrogen). Next, cDNA was synthesized from 1 μg of total RNA using a cDNA kit (Bioneer, Daejeon, Korea). Primers for genes related to inflammation were designed for PCR and qPCR. Reaction mixtures (20 μL total volume) contained 0.5 μM primer mixture, SYBR-Green with low ROX (BioFact, Daejeon, Korea), nuclease-free water (Ambion, Austin, TX, USA), and 2 μL of cDNA. Conditions were as follows: denaturation/activation for 10 min at 95°C, followed by 40 cycles of 15 sec at 95°C, 30 sec at 56°C, and 20 sec at 72°C. Reactions were performed on a StepOne Real-Time PCR instrument (Applied Biosystems, Foster City, CA, USA). Quantification of gene expression was based on the C_T_ value for each sample.

### ACE2 expression

Data regarding expression of ACE2 by human tissues and cell lines were obtained from The Human Protein Atlas database [47].

### Transfection and reporter assay

A 208-bp fragment of the 3’ UTR of the SARS-CoV-2 genome was synthesized by Bionics. The 3’ UTR fragment was digested with *Xba* I and inserted downstream of the firefly luciferase gene in pGL3-control (Promega, Madison, WI, USA), yielding pGL3covi-3UTR_Luc. All constructs were verified by sequencing. Human neuroblastoma SK-N-BE(2)C cells were maintained in DMEM supplemented with 10% FBS. All culture media contained 100 U/mL penicillin and 100 mg/mL streptomycin. For transfection, cells were plated at 1.2 × 10^5^ cells/well in 24-well plates in antibiotic-free DMEM 1 day prior to transfection. Transfections were carried out with Lipofectamine 2000 (Invitrogen). The total amount of DNA was 0.5 µg per well, comprising 0.2 µg of pGL3covi-3UTR_Luc and 0.3 µg of pRSV βgal as an internal control. Each transfection also contained the indicated miRNA. Twenty-four hours after transfection, cells from each well were lysed with 100 µL of lysis buffer (25 mM Tris-phosphate [pH 7.8], 2 mM DTT, 2 mM CDTA [1,2-diaminocyclohexane-N,N,N’,N’-tetraacetic acid], 10% glycerol, and 1% Triton X-100). An equal volume of firefly luciferase substrate was added, and luciferase activity was measured in a luminometer plate reader and normalized against beta-galactosidase activity.

### miRNA transfection

For miRNA transfection, 6 × 10^4^–2 × 10^5^ cells per well were seeded into a 24-well plate and transfected with 20 nM miRNA (hsa-miR-92a-3p, hsa-miR-26a-5p, hsa-miR-23a-3p, hsa-miR-103-3p, or hsa-miR-181-5p) (Bioneer) or with a Pre-miR miRNA Negative Control (Bioneer) using Lipofectamine 3000 (Invitrogen).

### RNA sequencing and data processing

Small RNA sequencing was performed using extracellular vesicles obtained under eight different conditions in pMSC media and two placenta tissue extracts. Sequencing was performed at the Beijing Genomics Institution (BGI, Shenzhen, China) using BGISeq-500. Reads were aligned to a human reference genome (GRCh38) using subread aligner [48] and the featureCounts tool was used to obtain miRNA read counts [49]. miRNA read counts were normalized, and miRNA expressed at low levels was filtered using the edgeR R package. Expression of each miRNA was transformed to CPM. Read quality control was performed using qrqc in R package and the distribution of small RNAs was calculated using sRNAtoolbox [50].

### miRNA target prediction and functional analysis

The PITA tool [51] was used to investigate miRNA binding sites in the 3’ UTR. The SARS-CoV-2 complete genome sequence (NC_045512.2) was obtained from the NCBI reference sequence database [52] and the 3’ UTR sequence was extracted from the SARS-CoV-2 complete genome. The PITA tool was used to predict binding sites for miRNAs. The default values were used for PITA analysis. Unconventional miRNA binding sites were predicted by the miRDB custom prediction tool [53]. To determine the biological function of miRNAs, experimentally validated targets were obtained from the miRTarBase [54]. miRNAs that made up a small percentage of the total reads (<1%) were excluded from the analysis. To investigate the biological function of miRNAs, GO term and KEGG pathway analysis were performed using the DAVID Bioinformatics Resource 6.8 [55]. The results of functional analyses were visualized using the Cytoscape 3.8 and Pathview R packages [56].

### Analysis of conserved miRNA binding sites

To investigate conserved regions within miRNA binding sites in SARS-CoV-2, the 3’ UTR sequence were obtained from the NCBI reference sequence database. The virus 3’ UTR sequence from the following coronaviruses were used: SARS coronavirus (NC_004718.3), SARS-CoV-2 (NC_045512.2), SARS coronavirus BJ01 (AY278488.2), Bat SARS coronavirus HKU3-1 (DQ022305.2), Bat SARS-like coronavirus SL-CoVZC45 (MG772933.1), Bat SARS-like coronavirus SL-CoVZXC21 (MG772934.1) and six complete genome of SARS-CoV-2 from Korean patients (MT730002, MT678839, MT304474, MT304475, MT304476 and MT039890). The 3’ UTR sequence of the coronaviruses were aligned using the MUSCLE tool [27]. The default value was used for MUSCLE analysis.

### Statistics

Statistical analysis was performed using either ANOVA, followed by Tukey’s HSD post-hoc test, or generalized linear model followed by a least square mean post-hoc (R basic functions and lsmeans package). For the qPCR results, all statistics were based on 2^(-delta Ct) values. A graph of qPCR results for inflammatory associated genes shows fold change values. A heatmap of the log2-CPM of miRNAs in pMSC-EVs was produced using gplots in the R package [57]. P values for GO term analysis and KEGG pathway analysis were corrected for multiple comparisons using the Benjamini–Hochberg method. Data are presented as the mean y standard error of the mean. A p value or adjusted p value <0.05 was considered significant

## Acknowledgments

This research was supported by the Bio & Medical Technology Development Program of the NRF funded by the Korean government, MSIP (NRF-2019M3A9H1103765)

---

**Figure S1:**
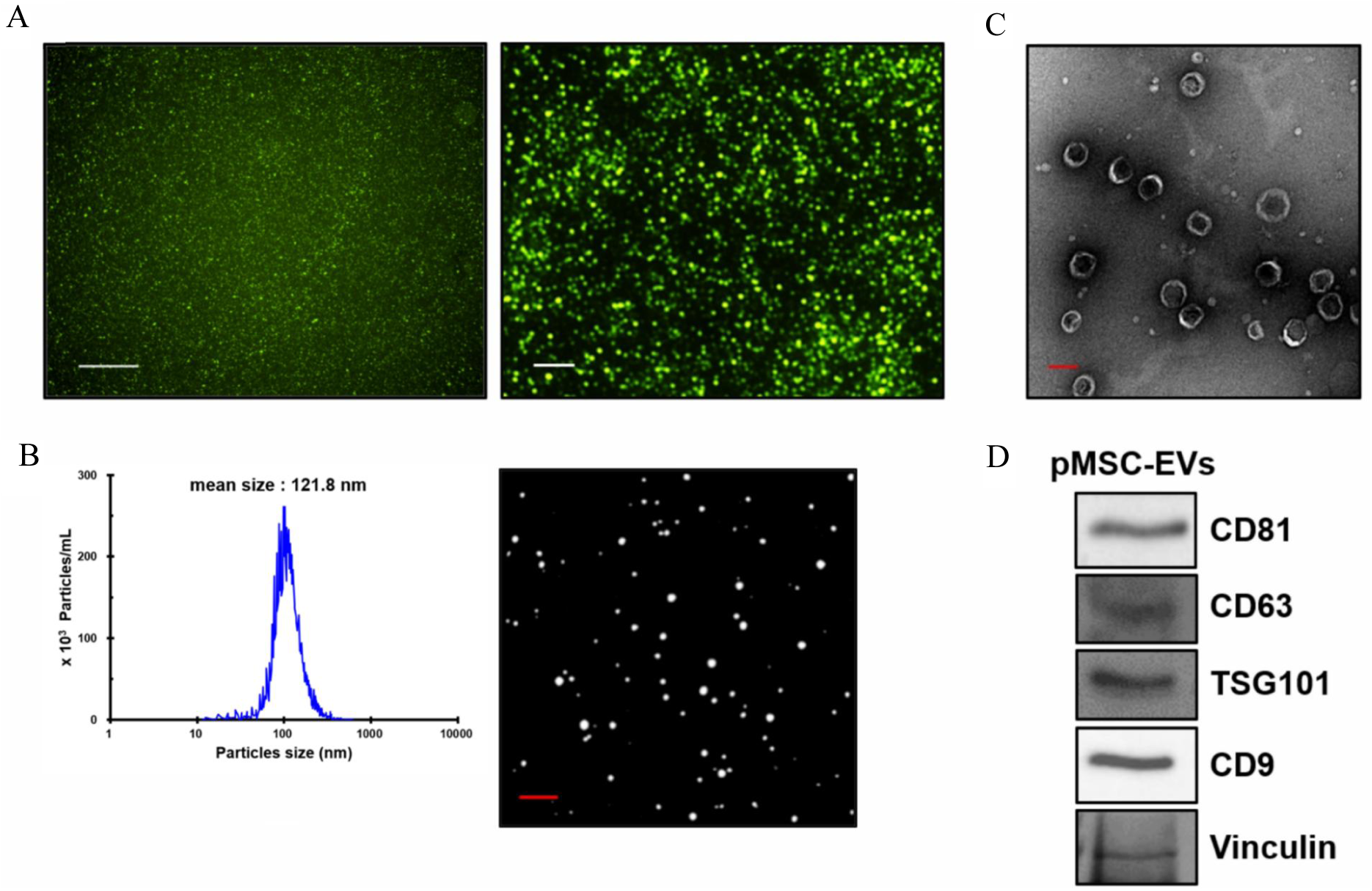
Characterization of human placenta mesenchymal stem cells-EVs. **(A)** EVs isolated from pMSCs were positive for the common EV marker CD63. Scale bar: 50 μm (Left; x40), 4 μm (right; x350). **(B)** The average size of EVs is 121.8 nm in diameter measured by nanoparticle tracking analysis (NTA) and **(C)** transmission electron microscopy (TEM). Scale bar: 0.1 μm. **(D)** Western blotting detected typical EV markers, including CD81, CD63, TSG101, and CD9.

**Figure S2:**
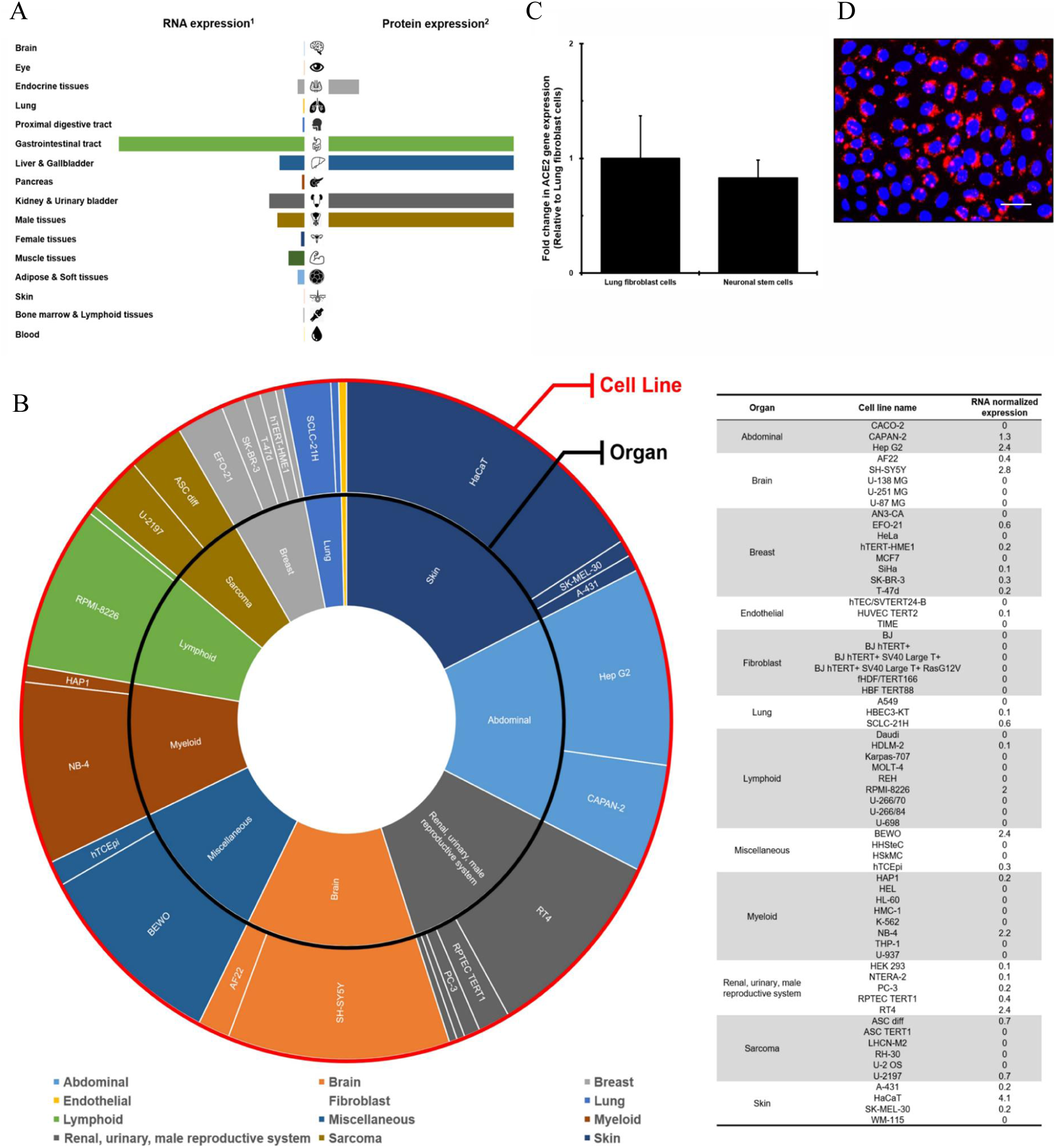
Organs expressing ACE2 receptors. **(A)** Summarization of ACE2 gene expression level of ACE2 receptor in various human tissues. Data were obtained from Human Protein Atlas (HPA).1 indicates ACE2 RNA expression measured by RNA sequencing.2 indicates ACE2 protein expression measured by *in situ* hybridization. **(B)** Summarization of ACE2 expression in human cell lines. Expression data were acquired from Human Protein Atlas (HPA). **(C)** ACE2 gene expression level of ACE2 receptor in human lung fibroblasts (LL24) and Brain stem cells (NSCs). **(D)** Uptake of EVs of PKH26 labelled EVs by Beas-2B cells observed for 24 h after EV treatment.

**Figure S3:**
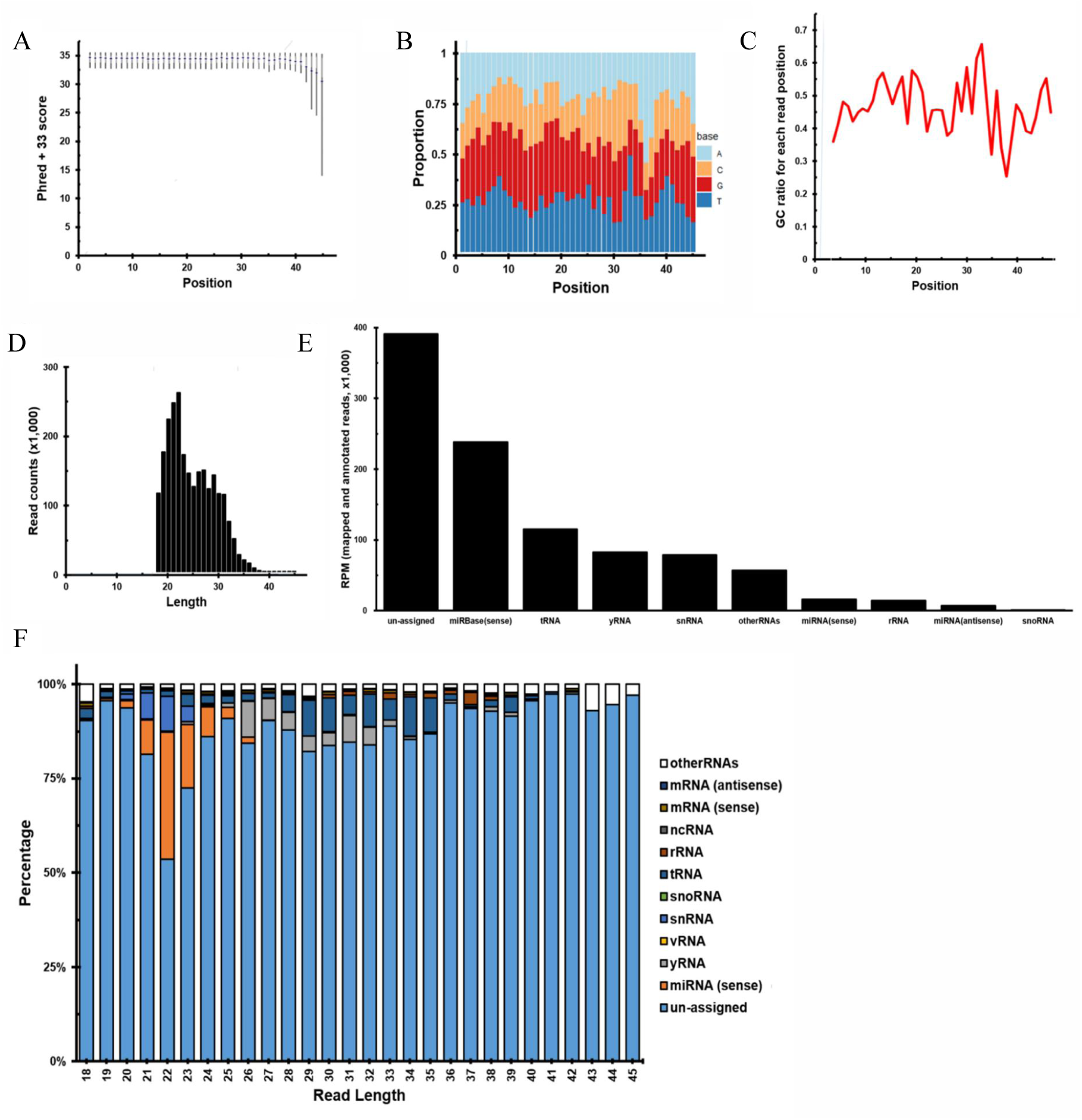
Quality control information for the small RNA sequencing library. **(A)** The Phred + 33 score for each read position. Total number of reads in the library, 24,834,367; average Phred score, 34.8. **(B)** The ratio of A, T, G, and C for each read position. **(C)** The GC ratio for each read position. **(D)** Histogram showing read length distribution. Read length ranged from 18 to 45. Most reads were 18–23 bases in length, corresponding to the size of miRNAs. **(E)** Table showing small miRNA library reads mapped on to the human genome. Unassigned reads were most common, followed by miRNAs, and tRNAs. **(F)** Classification of RNA versus read length. Reads of 20–24 bases, the size of most miRNAs, mapped mainly to miRNAs.

**Figure S4:**
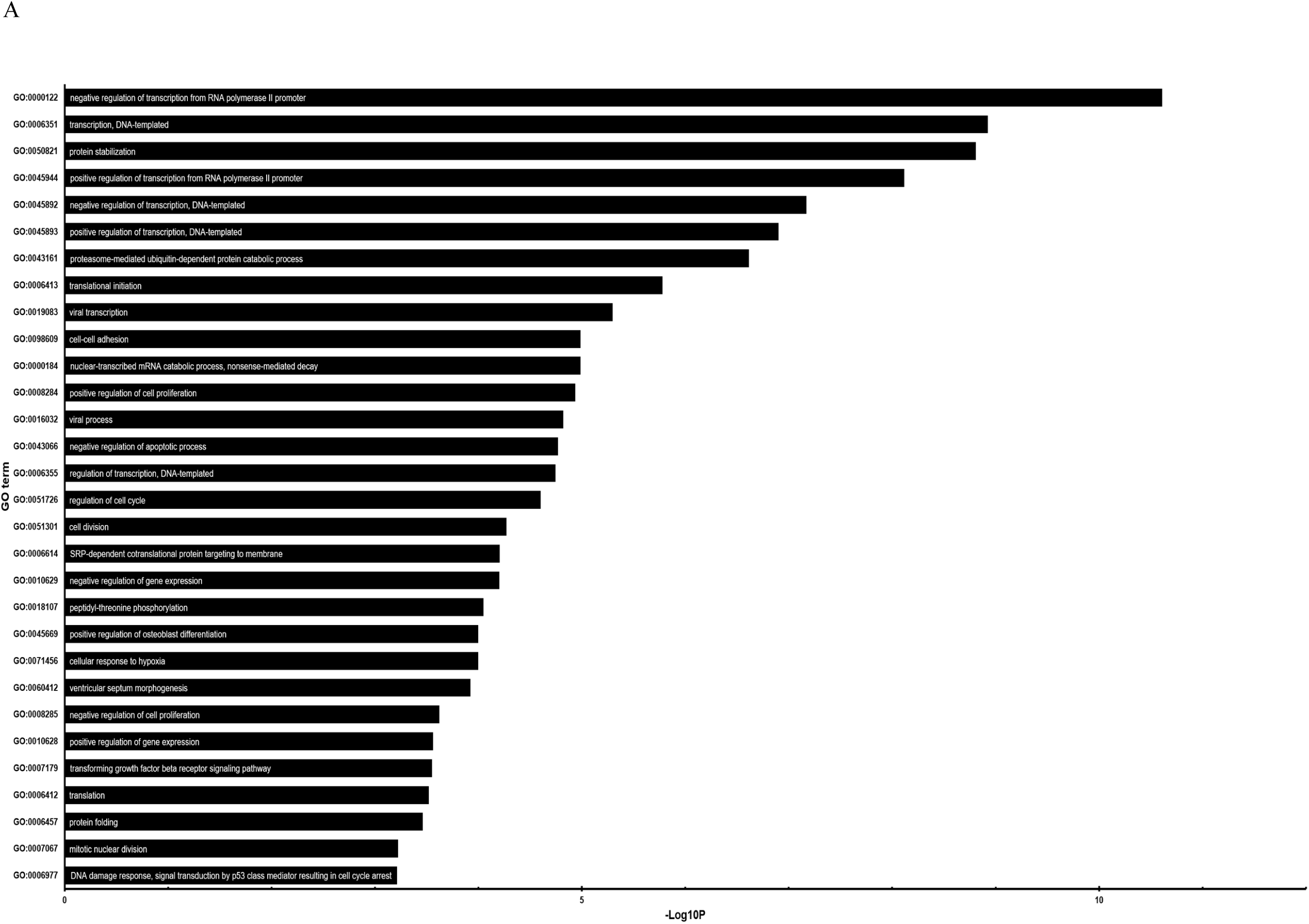
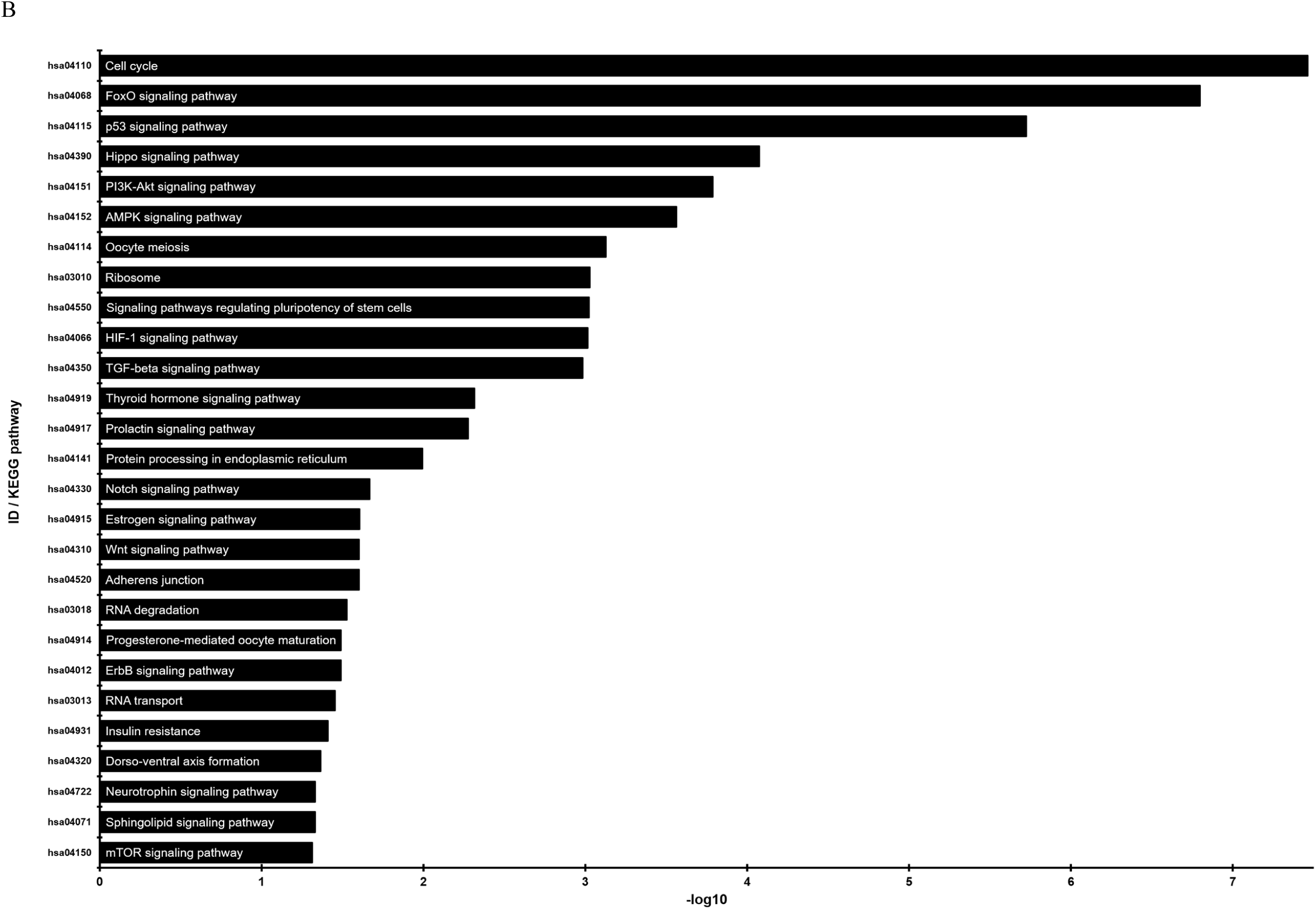
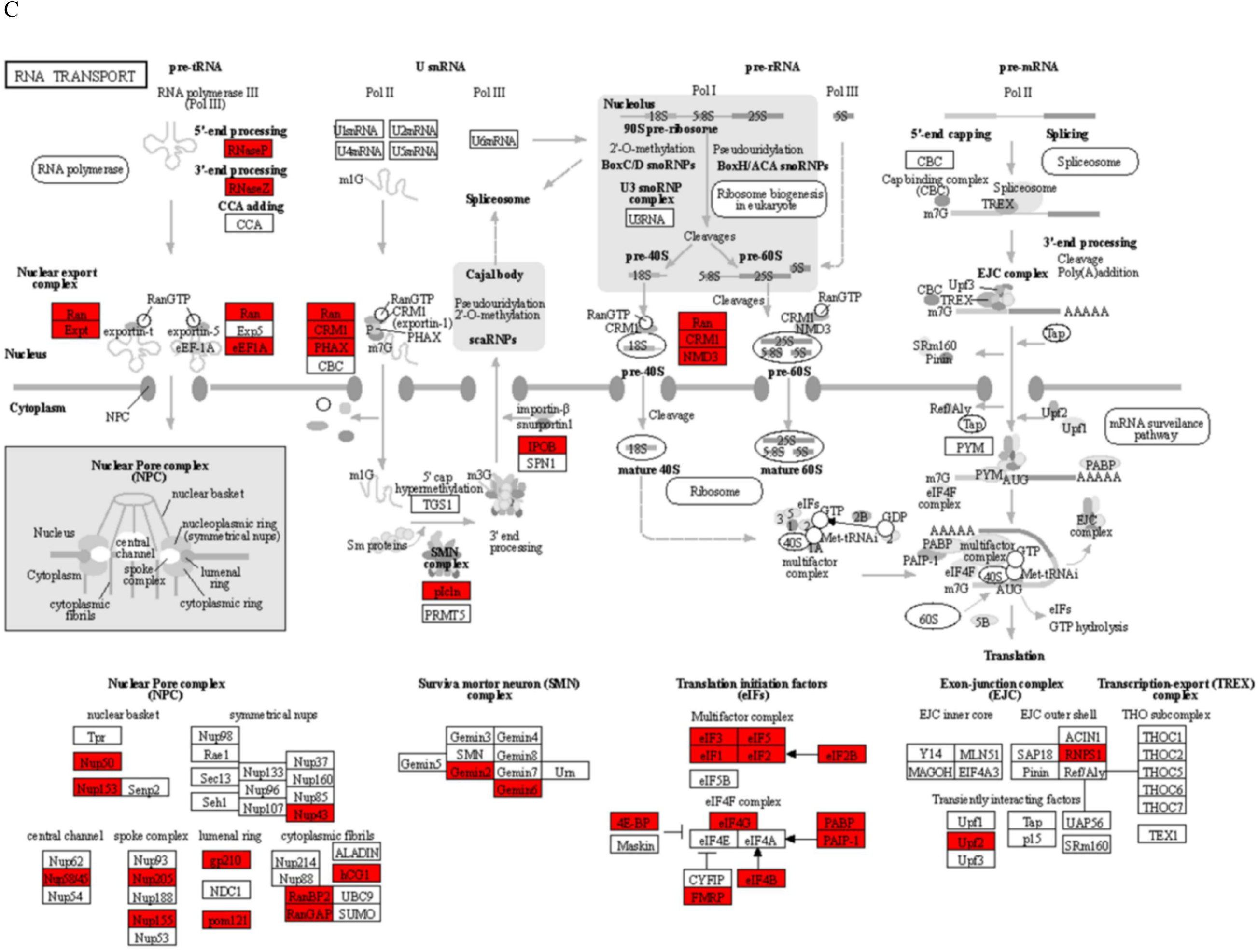
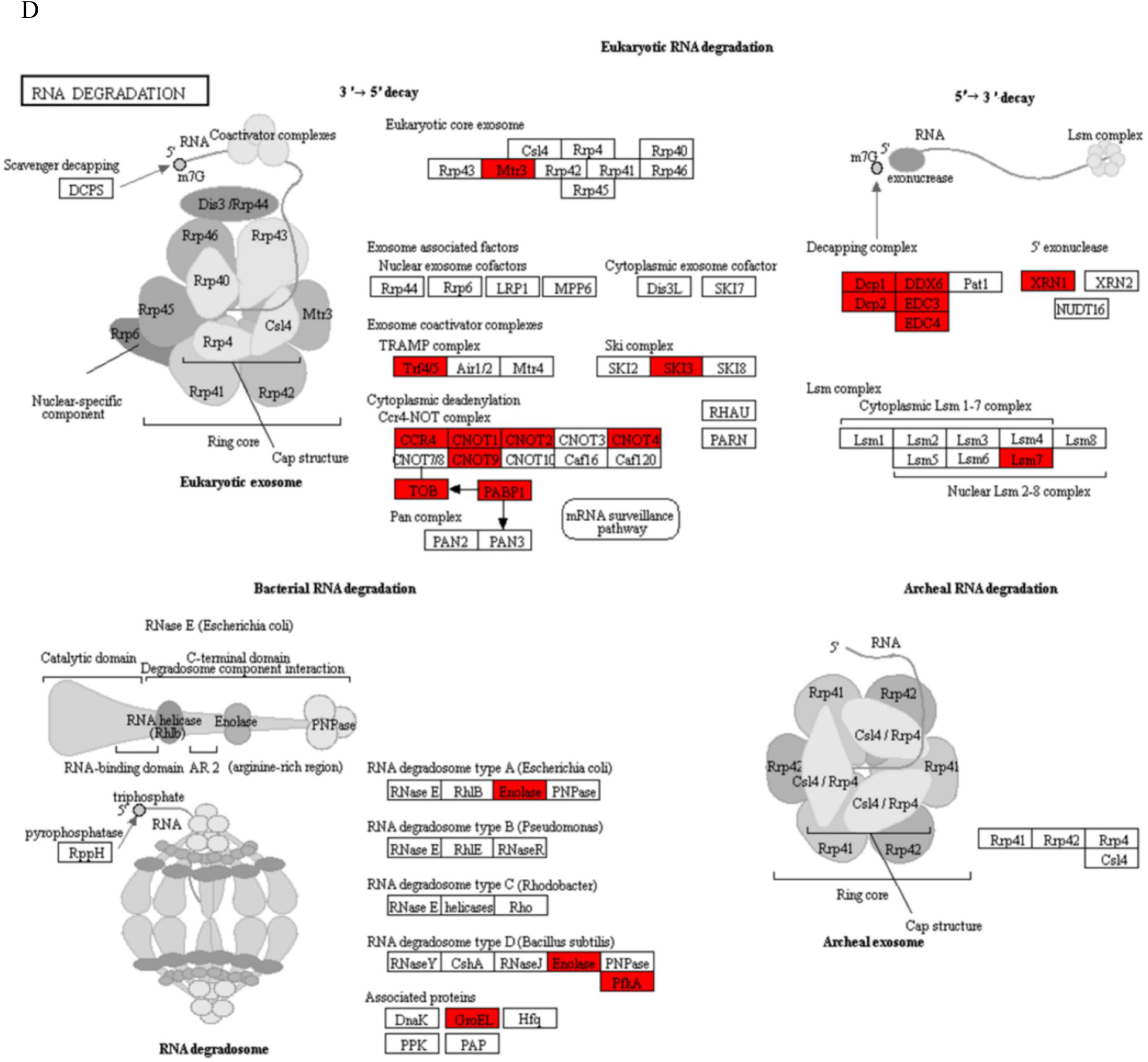
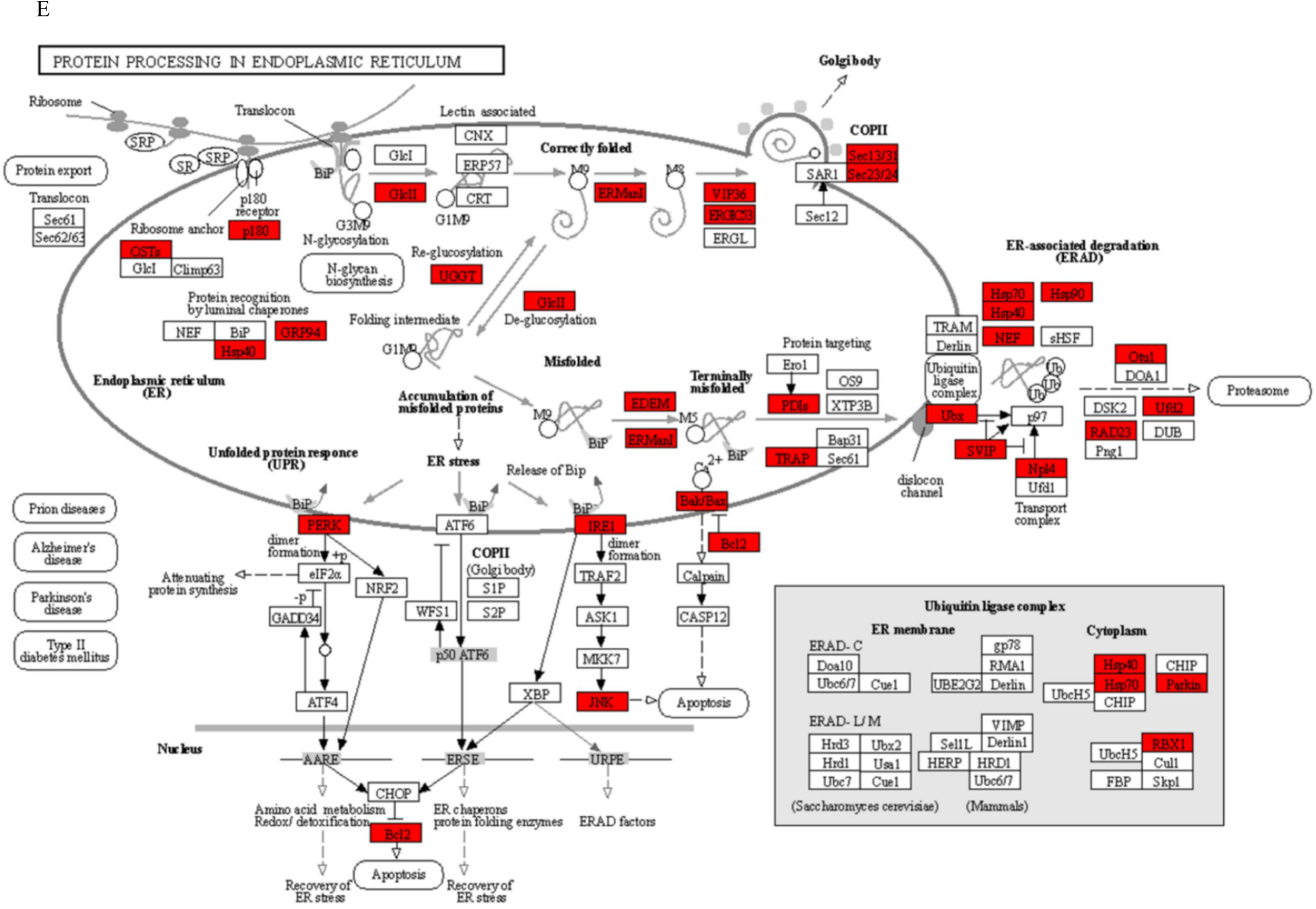
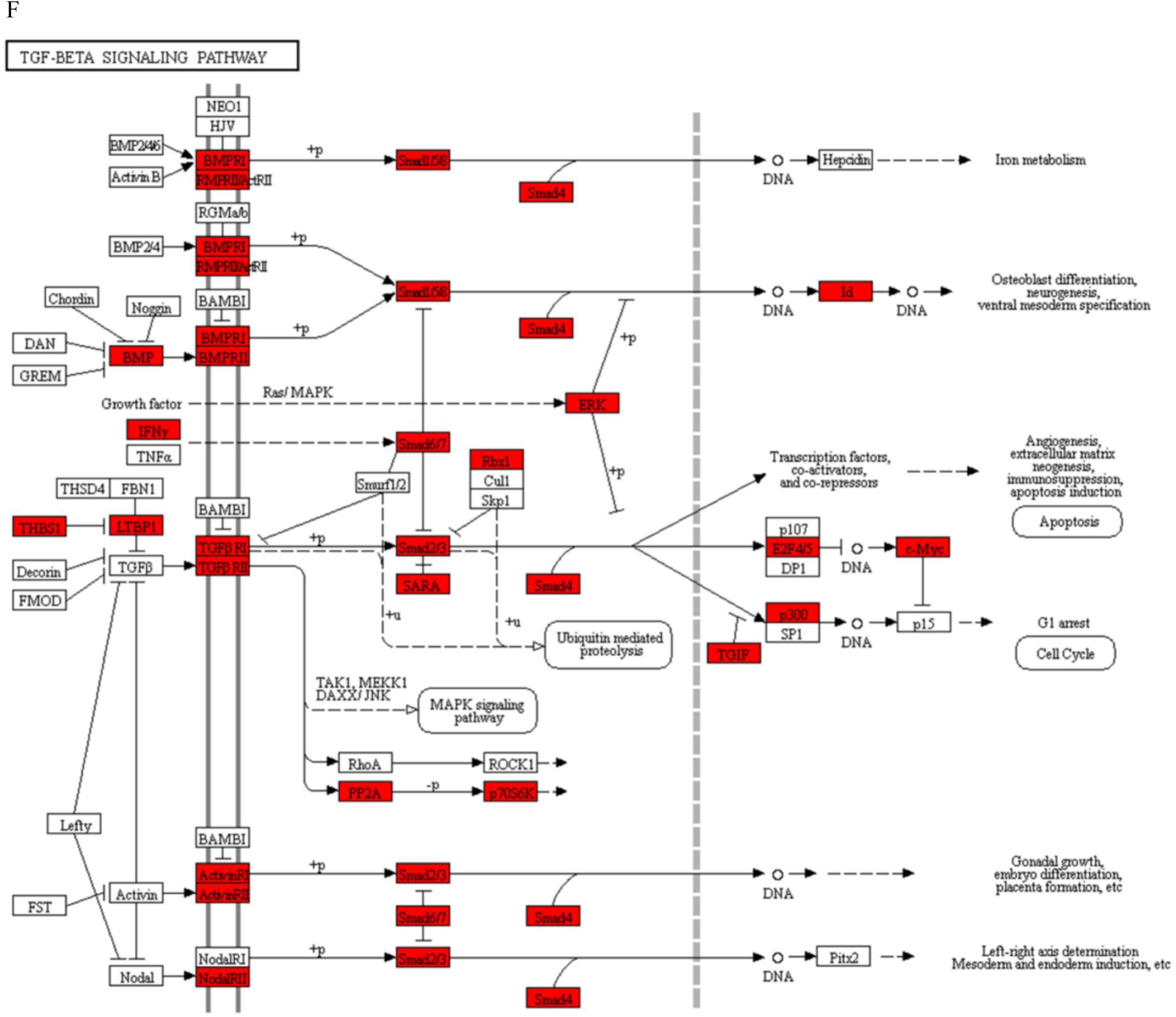
Function of mRNAs targeted by the five miRNAs **(A)** Statistical enrichment analysis of GO terms for the target genes of the miRNAs. The plot shows the top 30 GO terms. The x axis shows minus log10 transformed adjusted p-values and the y axis shows the GO terms. **(B)** Statistical enrichment analysis of 83 inflammatory associated genes in KEGG pathways. The pathway showing adjusted p-values <0.05 is presented in the plot. The x axis shows minus log10 transformed and adjusted p-values, and the y axis shows the names of the KEGG pathway i.e., RNA transport **(C)**, RNA degradation **(D)**, protein processing in endoplasmic reticulum **(E)**, and TGF-Beta signaling pathway **(F)**. The red box in each pathway indicates the gene targeted directly by miRNAs.

**Figure S5:**
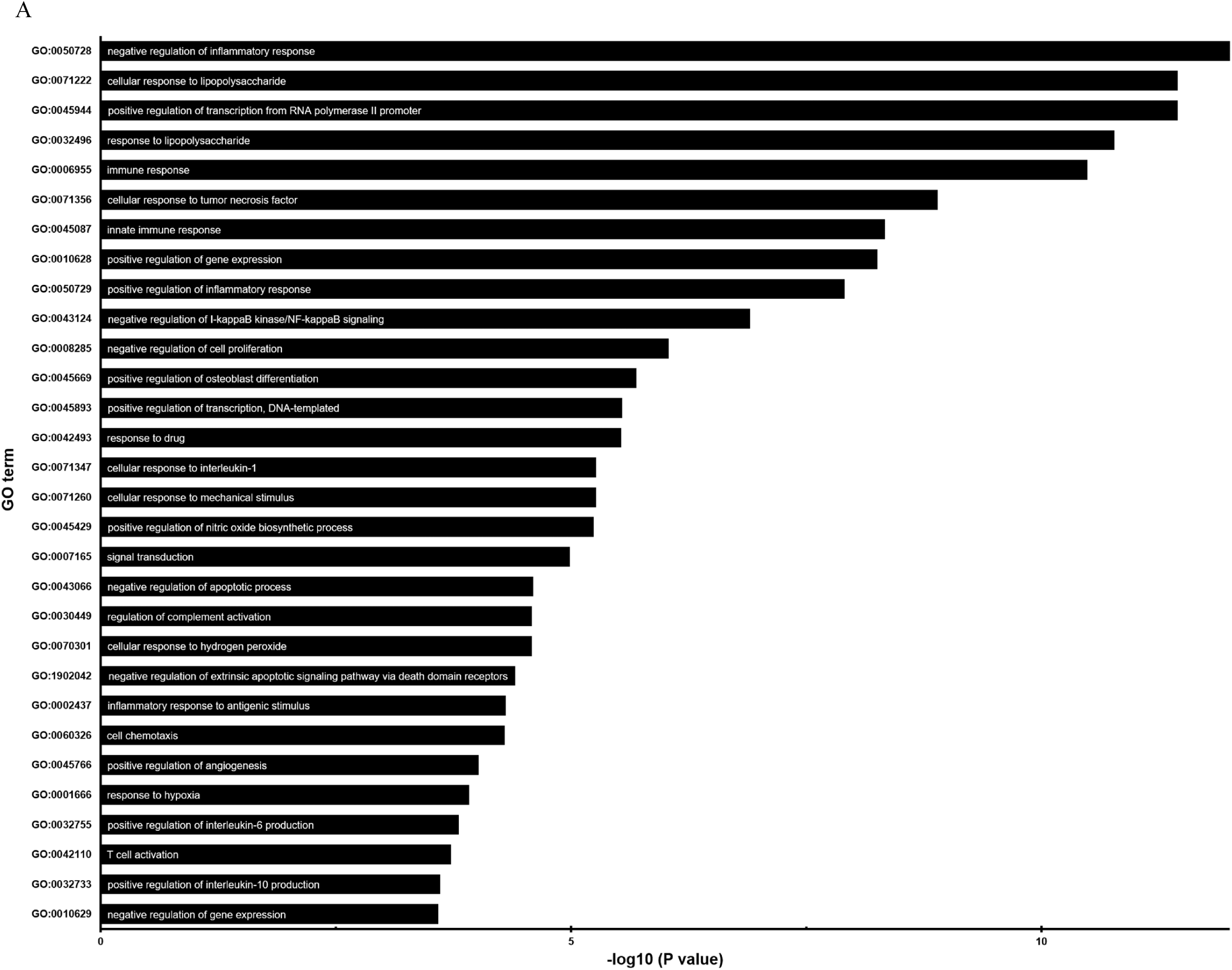
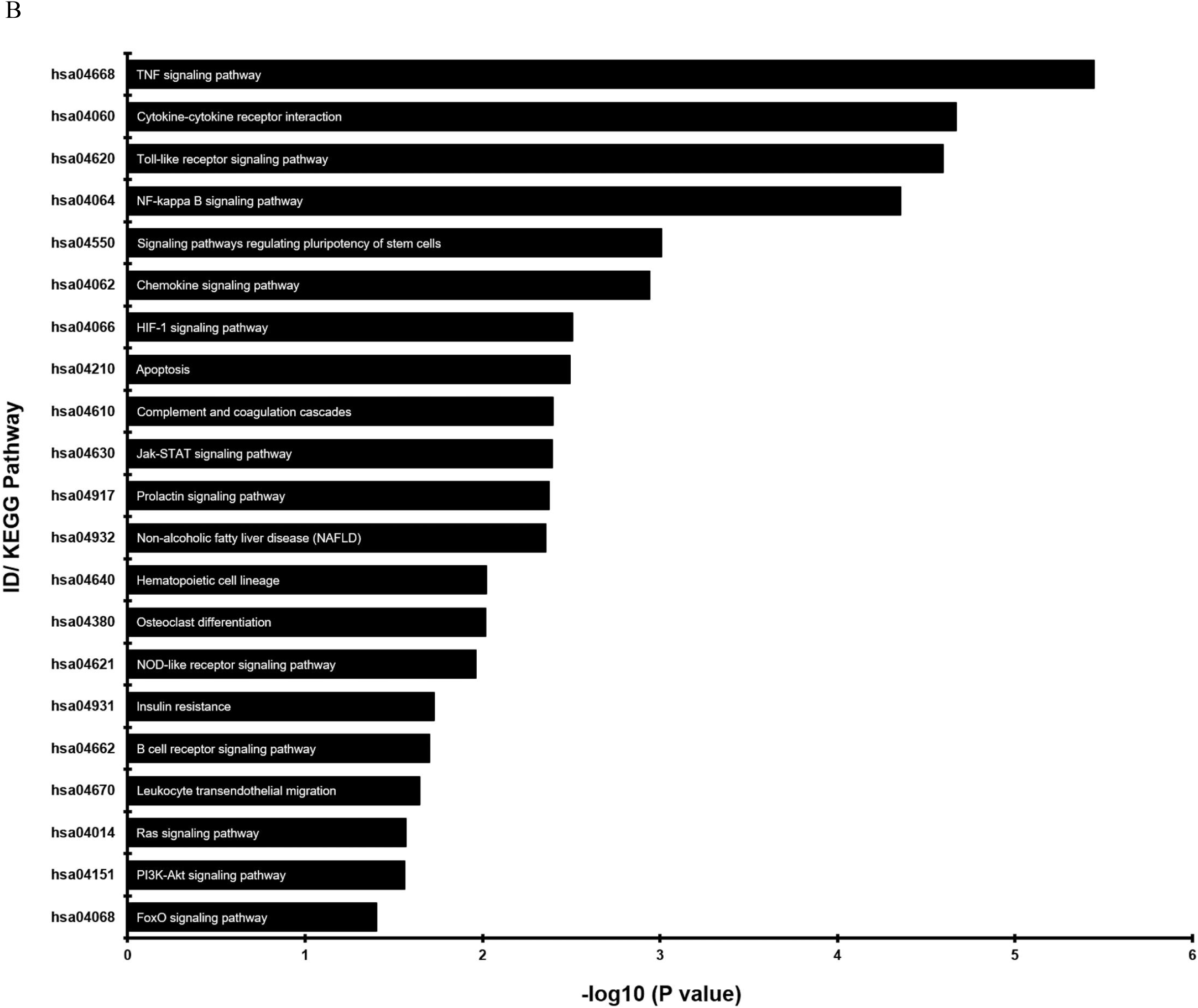
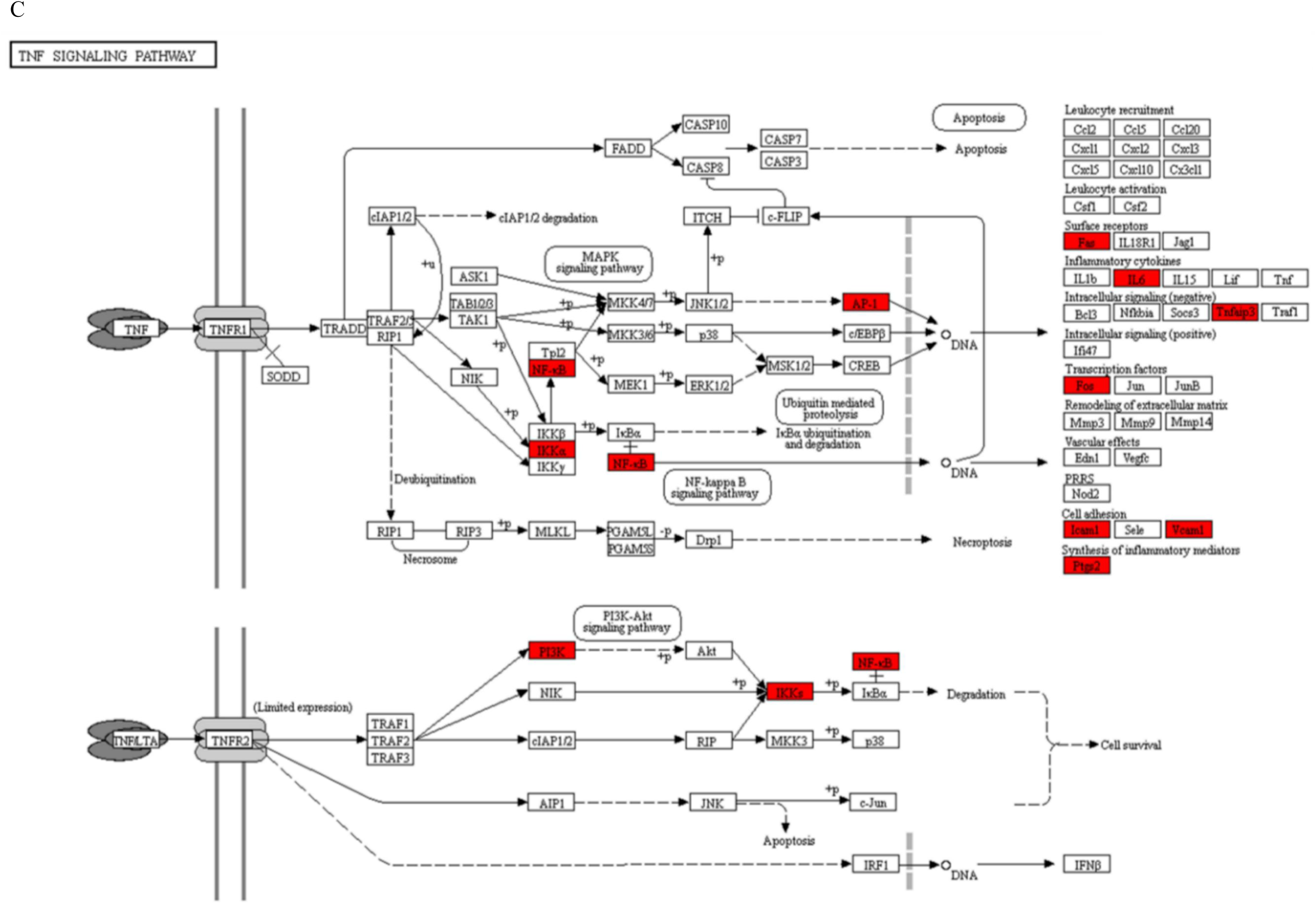
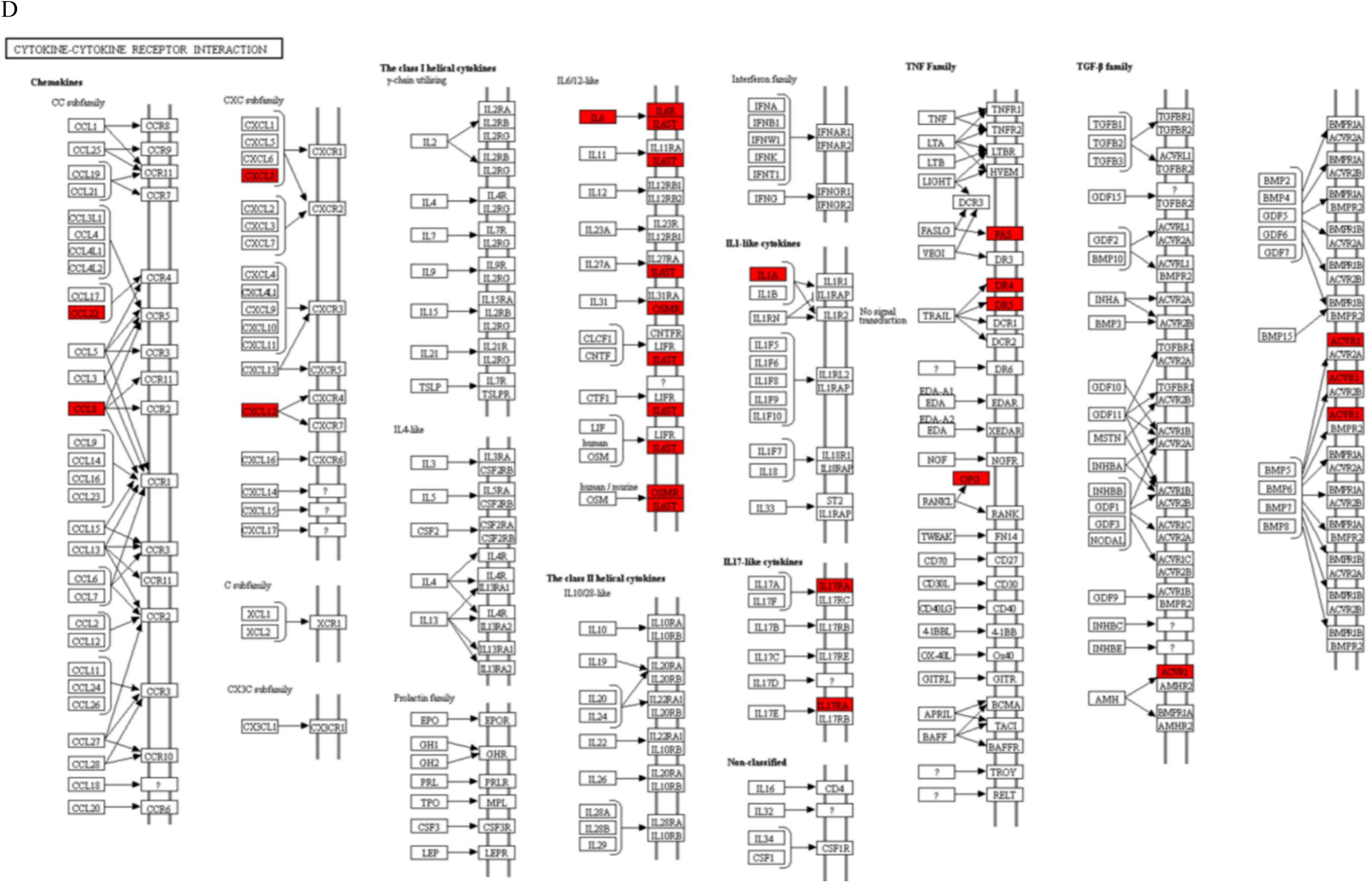
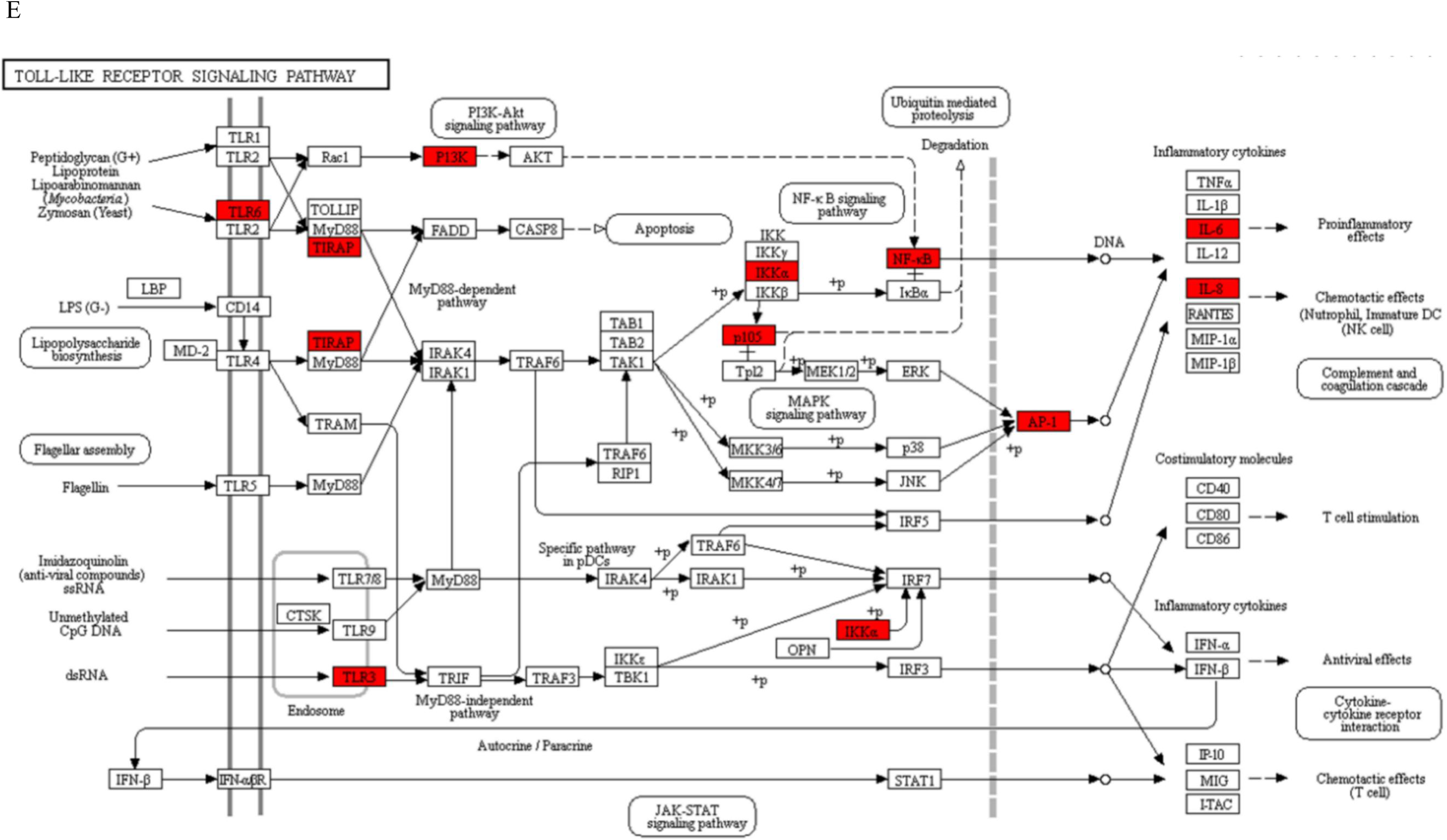
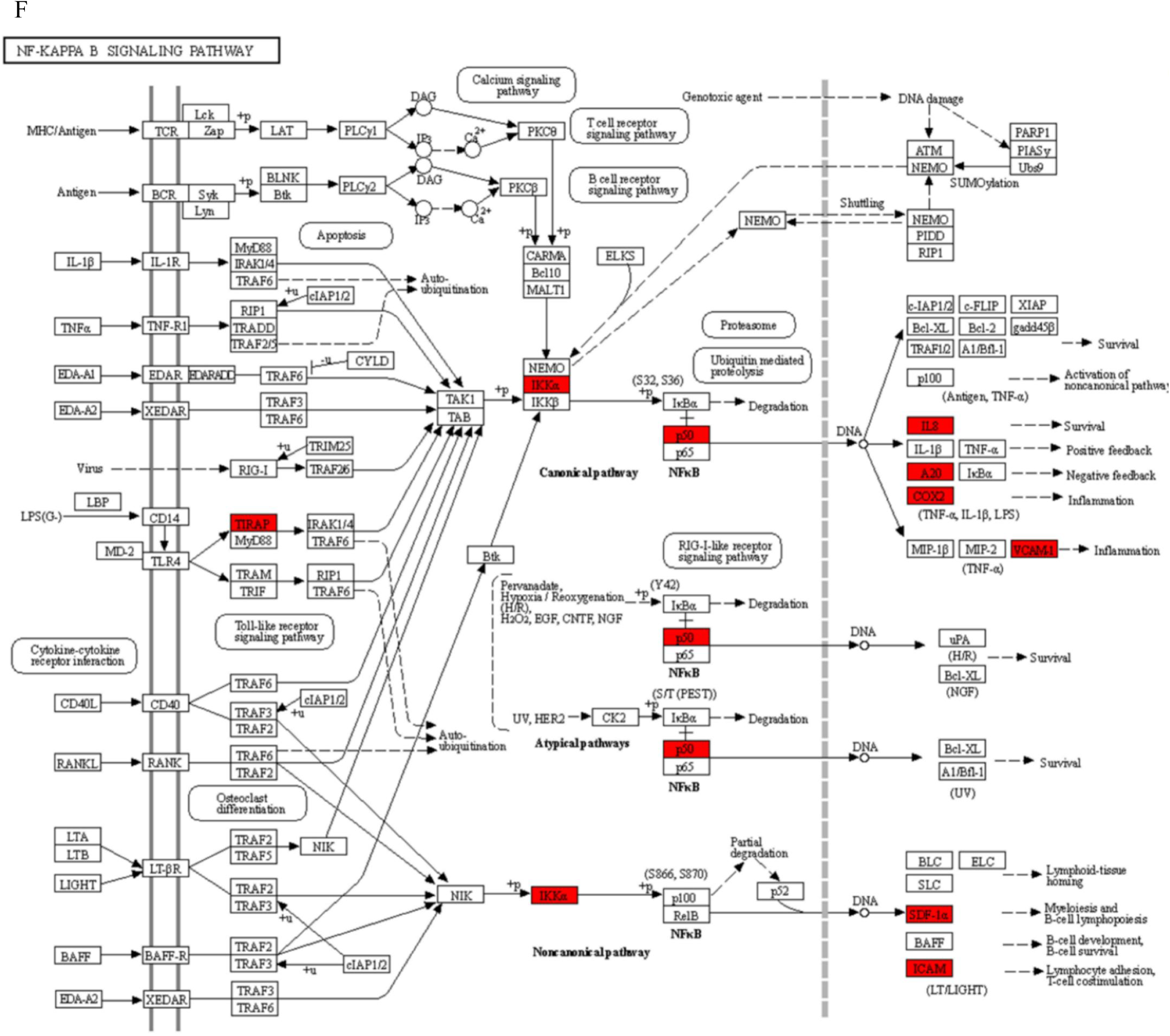
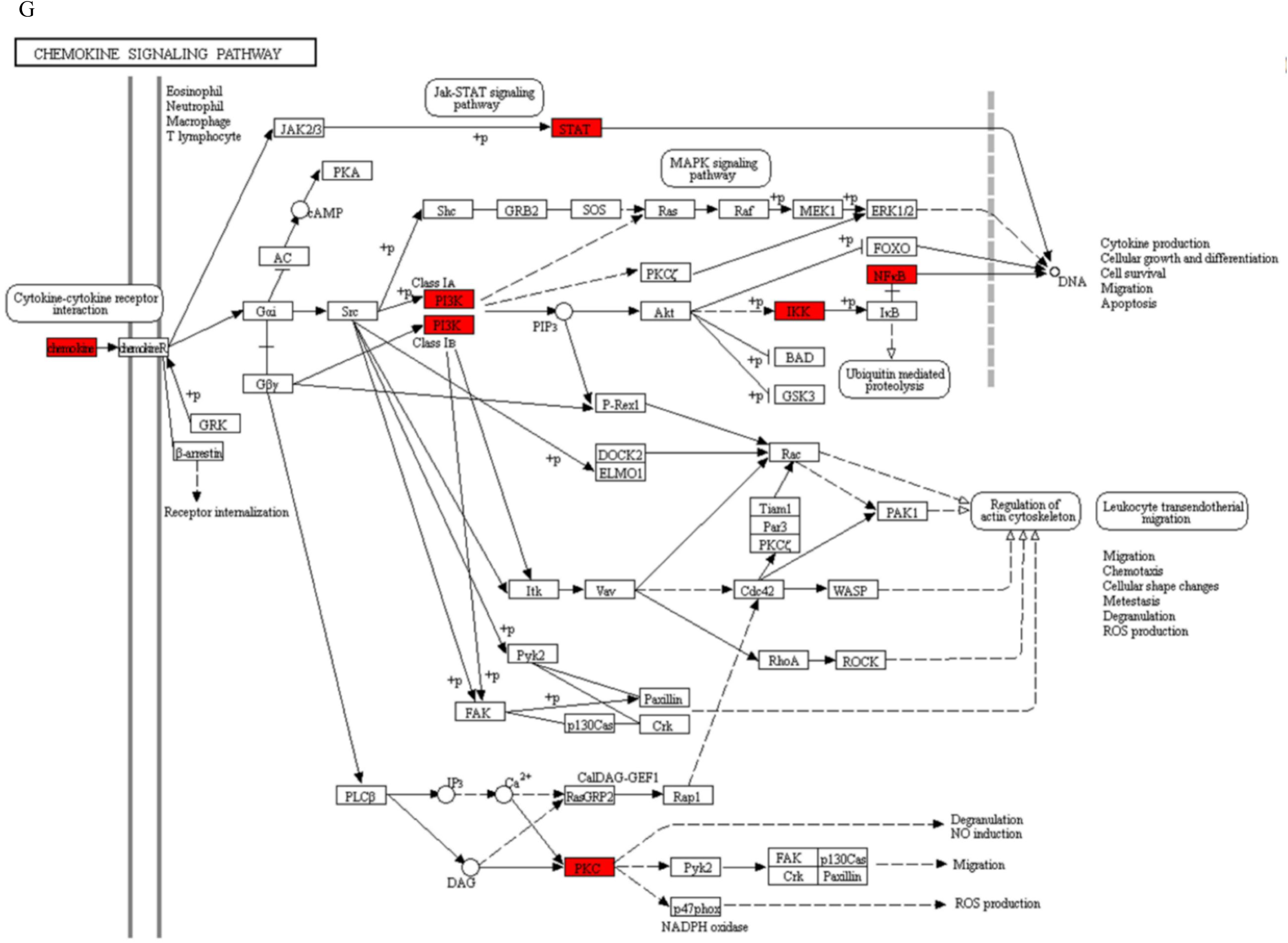
Significant inflammatory KEGG pathways regulating by the targets of the five miRNAs **(A)** Statistical enrichment analysis of GO terms for 83 inflammatory associated genes. The plot shows the top 30 GO terms. The x axis shows minus log10 transformed and adjusted p-values and the y axis shows the GO terms. **(B)** Significant pathways involving the 83 genes targeted by the five miRNAs. The 83 genes are predicted to be involved in inflammatory responses through pathways such as TNF signaling **(C)**, Toll-like receptor signaling **(D)**, cytokine-cytokine interactions **(E)**, NF-kappa B signaling **(F)**, and chemokine signaling **(G)**. The red box in each pathway indicates the gene targeted directly by the miRNAs.

**Table S1.**
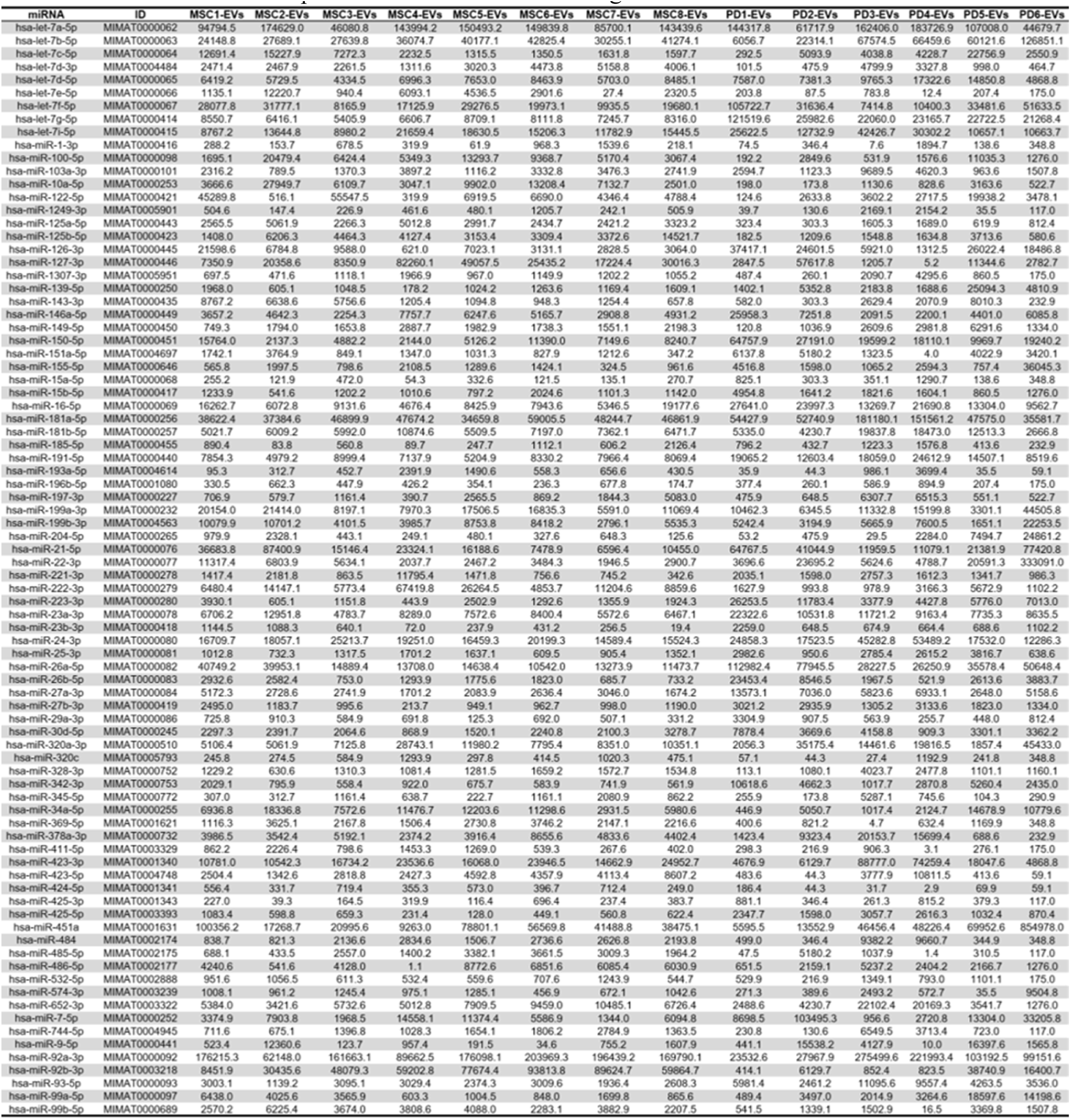
The expression levels of 84 common miRNAs present in EVs derived from eight MSCs cultured under various conditions and six PD-EVs.

**Table S2.** miRNAs matched to the coding region binding sites of SARS-CoV-2.

| miRNA | Expression (%)<sup>1</sup> | Score<sup>2</sup> |
| --- | --- | --- |
| hsa-miR-181a-5p | 4.69 | 70 |
| hsa-miR-26a-5p | 1.58 | 68 |
| hsa-miR-199a-3p | 1.31 | 69 |
| hsa-miR-320a-3p | 1.15 | 73 |
| hsa-miR-16-5p | 0.78 | 99 |
| hsa-miR-23a-3p | 0.74 | 79 |
| hsa-miR-181b-5p | 0.73 | 70 |
| hsa-miR-199b-3p | 0.65 | 69 |
| hsa-miR-122-5p | 0.64 | 67 |
| hsa-miR-378a-3p | 0.49 | 55 |
| hsa-miR-27a-3p | 0.29 | 67 |
| hsa-miR-103a-3p | 0.26 | 85 |
| hsa-miR-197-3p | 0.25 | 82 |
| hsa-miR-30d-5p | 0.20 | 95 |
| hsa-miR-26b-5p | 0.14 | 68 |
| hsa-miR-139-5p | 0.12 | 83 |
| hsa-miR-27b-3p | 0.11 | 67 |
| hsa-miR-15b-5p | 0.11 | 99 |
| hsa-miR-532-5p | 0.10 | 61 |
| hsa-miR-155-5p | 0.09 | 51 |
| hsa-miR-1-3p | 0.09 | 54 |
| hsa-miR-411-5p | 0.09 | 88 |
| hsa-miR-320c | 0.07 | 73 |
| hsa-miR-424-5p | 0.07 | 98 |
| hsa-miR-196b-5p | 0.06 | 52 |
| hsa-miR-29a-3p | 0.04 | 91 |
| hsa-miR-23b-3p | 0.03 | 79 |

<sup>1</sup> indicates “Percentage of miRNAs in EVs”
<sup>2</sup> indicates the prediction score or the miRDB method. According to miRDB, a predicted target with score > 80 is most likely to be real

